# A lipid metabolism defect is an underlying contributor to Diamond Blackfan anemia syndrome

**DOI:** 10.1101/2025.05.22.654334

**Authors:** Kaiwen Deng, Yu Wang, Joseph C. Min, Xiaofang Liu, Greggory Myers, Lei Yu, Susan Hammoud, Adham Adam, Rilie Saba, Vaneesha Natogi, Claire Drysdale, Brandon Chen, Hiroki Ueharu, Jennifer Lai Yee, Jacob Kitzman, Costas A. Lyssiotis, Yuji Mishina, Morgan Jones, Vesa Kaartinen, Yuanfang Guan, Rami Khoriaty, James Douglas Engel, Sharon A. Singh

**Affiliations:** Department of Computational Medicine & Bioinformatics, University of Michigan; Department of Cell and Developmental Biology, University of Michigan Medical School, Ann Arbor, MI; Department of Pediatrics, University of Michigan Medical School, Ann Arbor, MI; Department of Internal Medicine, University of Michigan Medical School, Ann Arbor, MI; Cellular and Molecular Biology Program, University of Michigan Medical School, Ann Arbor, MI; Department of Biologic and Materials Sciences & Prosthodontics, University of Michigan School of Dentistry, Ann Arbor, MI; Department of Human Genetics, University of Michigan Medical School, Ann Arbor, MI; Department of Molecular and Integrative Physiology, University of Michigan Medical School, Ann Arbor, MI; University of Michigan Rogel Cancer Center, Ann Arbor, MI

## Abstract

Analysis of neither Diamond Blackfan anemia syndrome (DBAS) cohorts nor animal models has revealed a potential mechanism for the variable anemia phenotype, a key feature of this disease. Here, we utilized an established *Rpl5^Skax23-Jus/+^* murine DBAS model in order to study this dynamic erythropoiesis deficiency. These haploinsufficient mice exhibit variably penetrant craniofacial and cardiac defects mimicking the phenotypes of DBAS patients bearing *RPL5* mutations. We additionally discovered that this specific heterozygous splicing mutation is pathogenic and leads to partial intron retention. By examining the transcriptome of fetal liver erythroid progenitors at E12.5, we demonstrate that the downregulation of erythroid differentiation pathways is consistent with the DBAS phenotype. We also identified dysregulated transcription of lipid metabolism genes with significant reduction in the abundance of *Scd1* in a subset of E12.5 mutant embryos at risk for erythroid failure. SCD1, a key enzyme that converts saturated to monounsaturated fatty acids, has not been previously linked to erythropoiesis or DBAS. When anemia was induced in adult mice, pretreatment with an SCD1 inhibitor resulted in improved erythropoiesis. This analysis suggests a key role of lipid metabolism in the variable anemia penetrance in DBAS and highlights a previously unappreciated pathway that may serve as a potential target for drug development.

**Key Points:**

- The variable anemia in *Rpl5^Skax23-Jus/+^*mice is triggered by intrinsic/extrinsic stress
- *Rpl5* haploinsufficient murine and human erythroid progenitors exhibit a lipid metabolism signature with downregulation of *Scd1*/*SCD*.

## Introduction

Diamond Blackfan anemia syndrome (DBAS) is a heterogeneous genetic disorder mainly caused by heterozygous ribosomal protein (RP) mutations^1,2^. This disorder was previously diagnosed in individuals with the classical presentation of macrocytic anemia in infancy, but with increasing recognition of variable phenotypes, the term DBA syndrome (DBAS) was more recently adopted. In addition to a defect in erythropoiesis, a subset of patients will present with defects in other hematopoietic lineages, leading to thrombocytopenia, neutropenia, or pancytopenia^3^. Some patients have birth defects, and there is an increased risk of developing cancer^4^. Developmental defects such as craniofacial malformations and cardiac abnormalities occur more frequently in *RPL* variants, especially *RPL5* (*uL18*) and *RPL11* (*uL5*), for unclear reasons^2,5,6^. Treatment-independence, previously termed remission, occurs in approximately 20% of individuals who previously required steroids or red cell transfusions. In some patients, the period of treatment-independence can be long, lasting for many years. Other patients develop periods of anemia cycling with treatment-independence but remain at risk for the development of malignancy^7^. The underlying mechanisms of this phenomenon remain unknown. We focused our studies on *RPL5,* as individuals with this genotype have very severe disease manifestations, specifically a higher rate of congenital anomalies, and a lower chance of developing treatment-independence^2^.

The initial discovery that most individuals with DBAS have a ribosomal protein mutation was intriguing and fostered increased interest in ribosome biology^8^. A key question that emerged was whether ribosomal protein haploinsufficiency leads to downstream developmental defects through reduced protein synthesis and/or possibly through extra-ribosomal functions of ribosomal proteins^9,10^. Many animal models and clinical cohorts have been examined to determine the underlying mechanism(s) and to develop new therapies^11^. These studies have demonstrated the role of nucleolar stress, heme toxicity, aberrant inflammatory pathways, and reduced translation of erythroid-specific factors as major drivers of erythropoiesis failure in this disorder^12^. Patients with severe anemia require red blood cell transfusions, but a subset of patients respond to long-term steroid treatment^7^. The finding of non-ribosomal mutations like *GATA1* and reduced GATA1 translation in DBAS has provided new avenues for broad therapies for patients, such as gene therapy^13–15^. However, despite tremendous progress, the mechanism for variable anemia among individuals with an identical *RPL5* genotype or within the same family remains unclear.

In our prior studies, we found a transient erythroid differentiation block at the CFU-E/proerythroblast stage in the E12.5 fetal livers of *Rpl5^Skax23-Jus/+^* mice, which began to resolve by E14.5^16^. Newborn pups had macrocytic anemia and increased mortality, but mice that survived weaning had no anemia or evidence of a hematopoietic stem/progenitor cell (HSPC) defect. We also demonstrated the variable presence of a ventricular septal defect (VSD) and a kinked tail phenotype with poor growth and increased mortality. Here, in order to investigate the mechanism that led to the erythroid differentiation defect, we examined in detail fetal liver erythroid progenitors using an unbiased whole transcriptome analytical approach. This strategy revealed dysregulation of lipid metabolism genes including downregulation of *Scd1* in mutant mice. Treatment of adult *Rpl5* mutant mice with a known SCD1 inhibitor did not induce anemia. Rather, SCD1 inhibition appeared to have a protective effect when anemia was induced with the hemolytic agent phenylhydrazine. We therefore propose that modulation of lipid metabolism and/or SCD1 may be a possible underlying mechanism leading to the variable anemia penetrance in DBAS. This warrants further study and may lead to the future development of therapeutics for DBAS.

## Methods

### Mice

We used *Rpl5^Skax23-Jus/+^* mice (MGI: 3046775), which we previously characterized^16^. For simplicity, we will refer to *Rpl5^Skax23-Jus/+^* mice as *Rpl5^+/-^*. Timed matings were performed to obtain E12.5 fetal liver (FL) cells and embryo sections were analyzed for the presence of VSD or craniofacial malformation as previously described^16^. CD71 positive/Ter119 negative cells were sorted to obtain early erythroid progenitor cells (BD FACS ARIA III or Sony MA900 Cell Sorter). Total RNA was extracted using RNeasy® Micro Kit (Qiagen, Cat 74004) and used for bulk RNA-seq analysis. Analysis was performed using DESeq2 and GO enrichment analysis. Additional details provided in supplemental methods, Tables S1 and S2.

### RNA-seq data processing

Both library preps, next generation sequencing runs, and data pre-processing were conducted by the University of Michigan Advanced Genomics Core. Paired-end 151 bp raw reads retrieved from the experiments were trimmed using Cutadapt v2.3^24^ and mapped to the GRCm38 assembly (ENSEMBLE) using STAR v2.7.8a^22^. Gene counts from the alignments were estimated using RSEM v1.3.3^25^. Alignment options followed ENCODE standards for RNA-seq. Quality controls were reported by FastQC v0.11.8^26^, Fastq Screen v0.14^27^, and Multiqc v1.7^28^. Gene information was re-annotated with AnnotationDbi^29^ and *org.Mm.eg.db* to retrieve the Entrez Gene identifiers (Entrez ID) and gene names based on their ENSEMBL IDs.

### Statistics

GraphPad Prism 10 was used to graph data and conduct the Student’s unpaired t-test to determine statistical significance. Error bars represent the standard deviation from the mean. (* p<0.05; ** p<0.01, *** p<0.001, **** p<0.0001 for all figures).

## Results

### *Rpl5^Skax23-Jus/+^* mice have evidence of a cleft palate

We previously described *Rpl5^Skax23-Jus/+^* mice, with a heterozygous intronic *Rpl5* mutation that leads to *Rpl5* haploinsufficiency^16^. Mutant mice are small with partially penetrant kinked tails, macrocytic anemia, and a ventricular septal defect but no craniofacial defect at birth. The severe phenotype in this model aligns closely to human DBAS patients with *RPL5* mutations^2^. However, it was unclear if these mutant mice developed craniofacial defects, which is a major phenotype of *RPL5* mutations in humans^5^. Further embryo characterization at E15.5 revealed that 20% of *Rpl5^+/-^* mice had a VSD (Table S3) and 30% of *Rpl5^+/-^* mice showed lack of palatal fusion (Figure 1A, Table S3). The lack of cleft palate and/or VSD in adult animals indicates that severe morphological defects might only be associated with early embryonic/perinatal mortality. Additional histological observation of the facial structures by hematoxylin/Alcian blue was performed at E14.5, which again demonstrated defects in palate elevation and tongue descent (Figure 1B). We performed staining for proliferation (pH3) and cell death (TUNEL) at E14.5 (Figure 1C). The mutant mice had significantly higher pH3-positive cells but a trend towards lower numbers of TUNEL-positive cells compared to WT (Figure 1D). This indicates dysregulation of normal patterns of cell growth and apoptosis during craniofacial embryogenesis caused by *Rpl5* haploinsufficiency.

**Figure 1.**
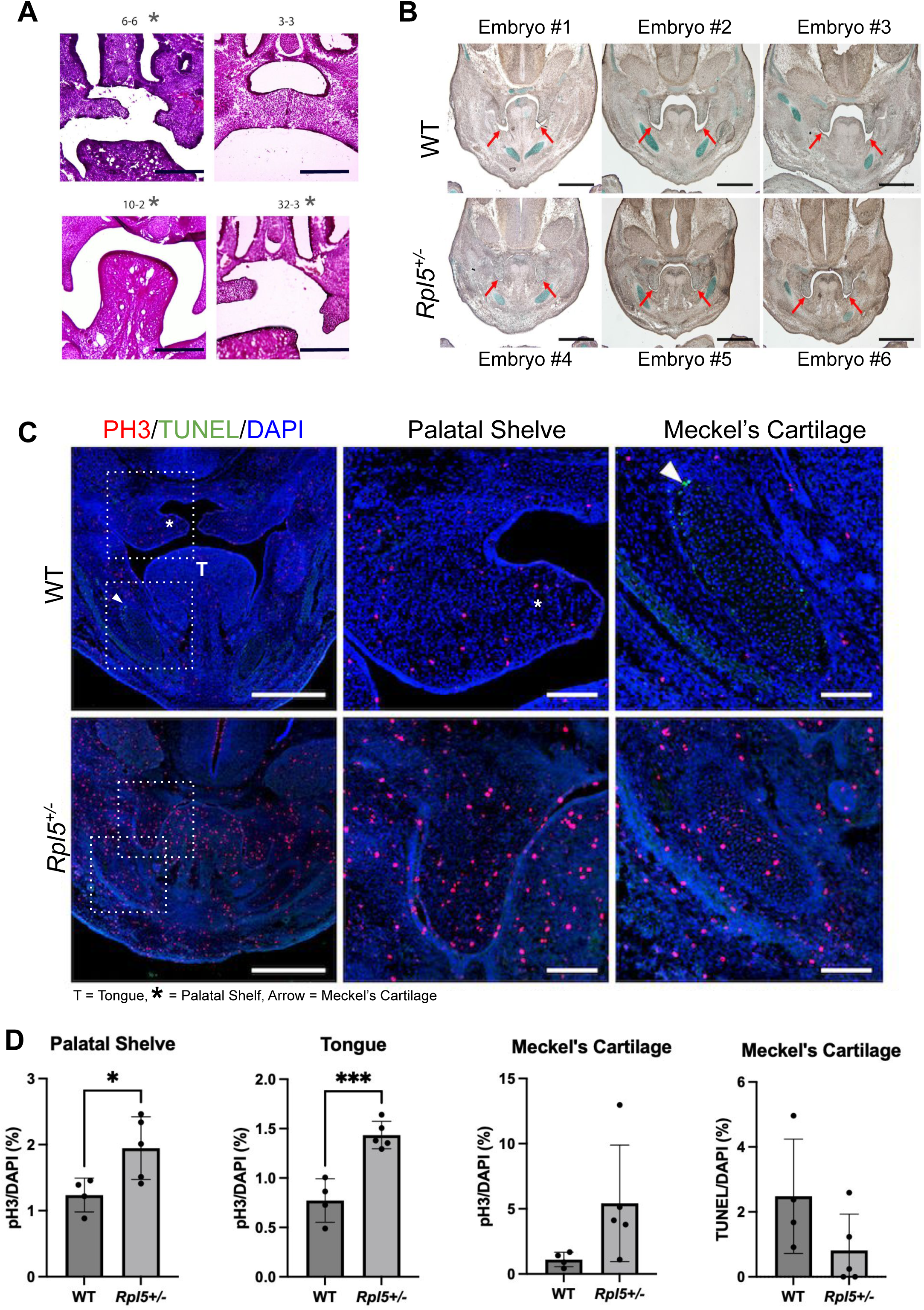
*Rpl5^Skax23-Jus/+^* mice have a cleft palate during embryonic development. (A) Histological characterization of four litters at E15.5 (WT, n=11; *Rpl5^+/-^*, n=13) with representative histology sections from 4 mutant mice. Mutant #3-3 had no cleft palate while the other mutants demonstrated lack of palatal elevation and/or fusion (scale bar: 300µm). (B) Analysis of E14.5 embryos (WT, n=3; *Rpl5^+/-^* n=3) after Alcian blue staining showed a delay in normal palate elevation in mutant mice. (C) Further staining (WT, n=4; *Rpl5^+/-^* n=5) indicated an increase in proliferation (pH3) and a decrease in apoptosis (TUNEL) in *Rpl5^+/-^*cells (2 representative images presented). Scale: 500 µm and 100 µm (enlarged area). (D) Quantification of PH3 and TUNEL in palatal shelf, tongue, and Meckel’s cartilage.

### The c.3+6T>C intronic mutation results in alternative splicing

Our prior work demonstrated that the ENU-induced *Rpl5* mutation (c.3+6T>C) was at a highly conserved intronic site and resulted in *Rpl5* haploinsufficiency^16^. Concordantly, RNA-seq of E12.5 CD71+ Ter119-fetal liver cells in this work also showed a shift away from the canonical isoform in mutant cells (Figure 2A). To test this splicing disruption as the underlying mechanism, we performed a minigene assay using the wild-type and mutant alleles of the *Rpl5* first exon. After transfection in HEK-293T cells and RT-PCR, we analyzed the spliced minigene transcript product by gel and next-generation sequencing (Figures 2B, C). In the WT construct, the splice donor immediately following the start codon was predominantly used, resulting in a majority of in-frame transcripts (∼92%). By contrast, in the mutant construct, usage of this donor was almost entirely abolished (∼0%).

**Figure 2.**
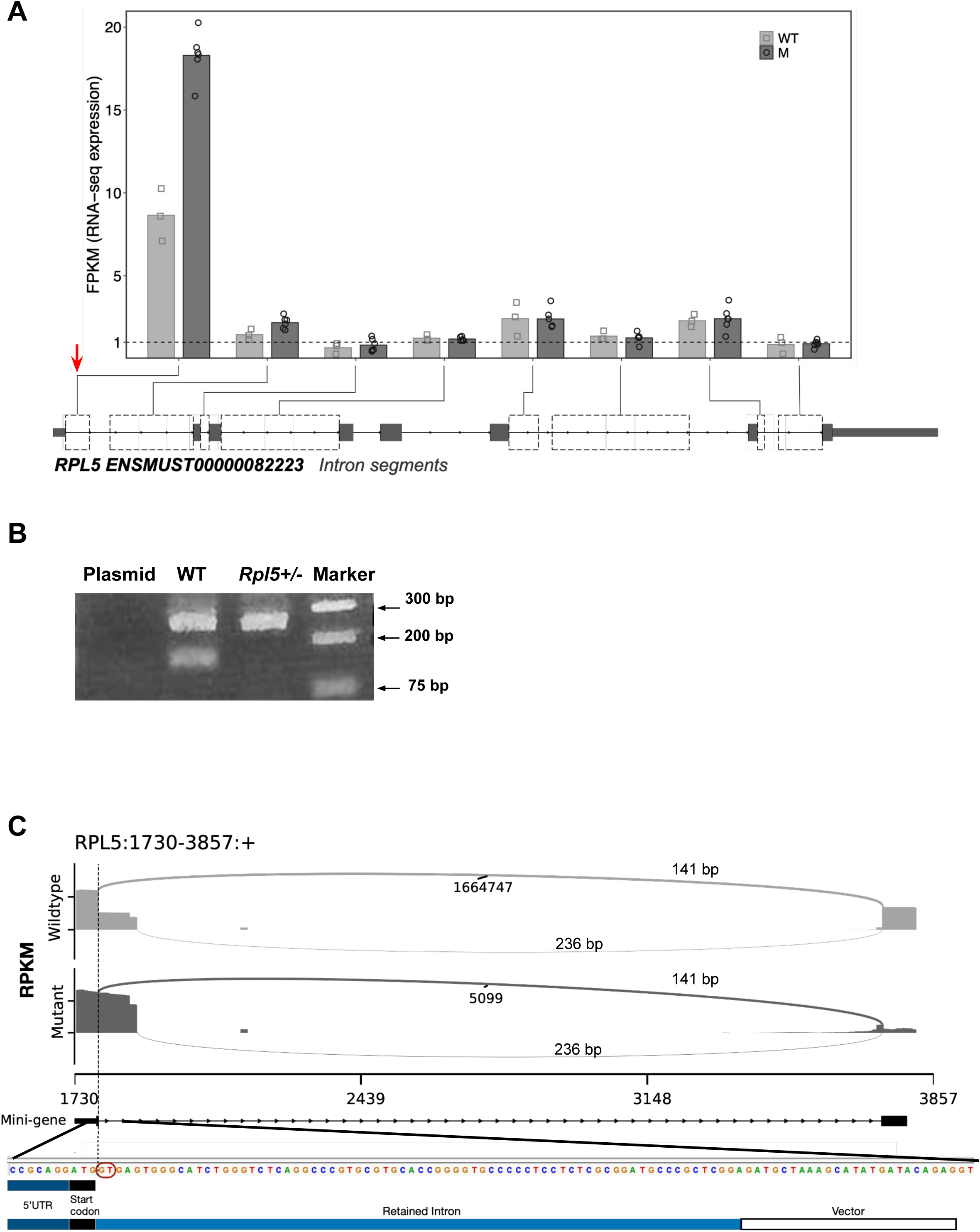
The mechanism of *Rpl*5 haploinsufficiency. (A) Bioinformatic analysis of RNA-seq data shows partial intron retention caused by the mutation in the first intron segment of *Rpl5*.The arrow indicates c.3+6T>C intronic mutation (B) Gel electrophoresis of the nested PCR product from the minigene assay showed a predominantly larger-size PCR product of the mutant compared to the WT sequence (WT, two bands 141bp, 236bp; M (*Rpl5^+/-^*), 236bp): lower band (141bp) only seen in WT due to abolished canonical splice donor in mutant sequence. The upper 236 bp band is created by synthetic vector splice site and is used by both WT and mutant. (C) NGS analysis from the minigene assay showed a disruptive splicing pattern with the *Rpl5* c.3+6T>C intronic mutation compared to the WT. The sashimi plot revealed the spliced reads in the mutant versus the WT. The y-axis is the normalized reads in RPKM. The vertical dashed line indicates the cutting site after the first exon. The numbers without “bp” are the junction coverages in raw read counts, and the numbers with “bp” are the lengths of the junctions.

### E12.5 *Rpl5^+/-^* fetal liver erythroid progenitors have reduced erythroid differentiation and lipid metabolism gene expression

In these mice, we previously reported a severe erythroid differentiation block at E12.5. To further investigate the cellular mechanism of this defect, we sorted E12.5 erythroid progenitors in the CD71+ Ter119-gate and performed mRNA-seq analysis (Figure 3A). Due to the extremely low cellularity of this progenitor cell population in mutant animals, we used pooled embryos to generate WT and mutant samples. The differential expression (DE) analysis identified 220 highly variable genes with adjusted p-values ≤ 0.005 and absolute log2 fold changes (LFC) ≥ 0.99 (Figure 3B), among which 142 of the genes were down-regulated in the mutant, and 78 were up-regulated (Figure 3C). Gene ontology (GO) enrichment analysis highlighted 271 significant terms from all variable genes (q-values ≤ 0.05) and 52 from the down-regulated ones (Figure S1). Based on these top significant terms and our analyses of previously published human data^39^ (Figures S2, S3), we selected erythrocyte-related, lipid-related, and oxidative-stress-related terms for further investigation. Nine erythrocyte and lipid related terms demonstrated significant enrichment among all significant and down-regulated genes (Figure 3D). Analysis from gene set enrichment analysis (GSEA) using all expressed genes further supports the observations related to erythropoiesis (Figure 3E). We found seven down-regulated genes involved in erythrocyte development, differentiation, and homeostasis networks, consistent with the erythroid differentiation defect we previously described (Figure 3F). We also found two up-regulated and six down-regulated genes involved in several lipid metabolism networks, including the neutral lipid, glycerolipid, triglyceride, and acylglycerol biosynthetic/biosynthetic processes (Figure 3G).

**Figure 3.**
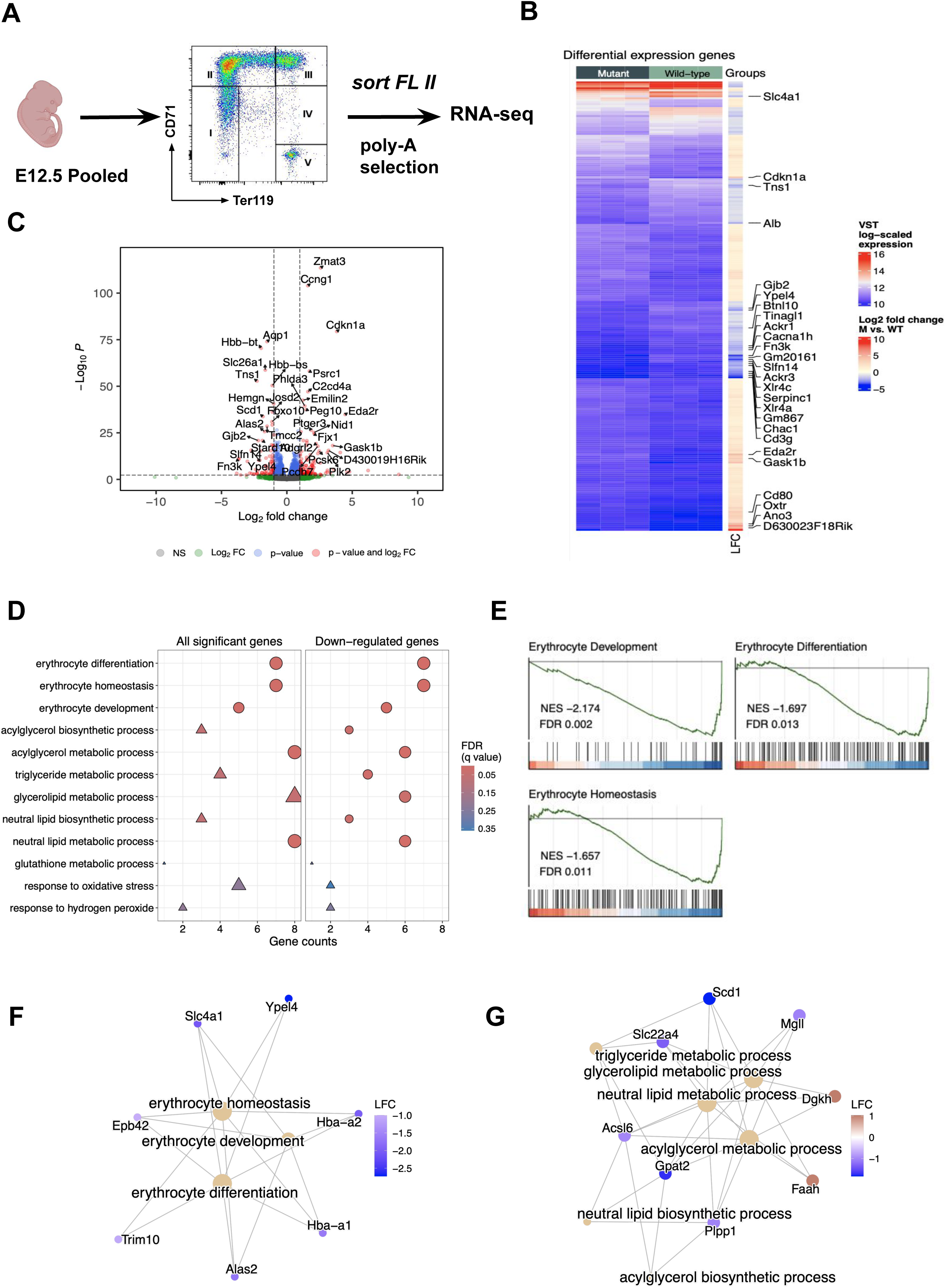
Transcriptomic analysis of E12.5 fetal liver (FL) erythroid progenitors. (A) We sorted E12.5 FL and performed bulk RNA-seq using mRNA from CD71+ Ter119-cells (WT, n=3; *Rpl5^+/-^* n=3 using pooled embryos) (II) (B) We performed differential gene expression analysis and data visualization via heatmap with the variance stabilizing transformation (VST) log-scaled expressions for all the significant genes. The rows and columns of the heatmap are automatically clustered. The corresponding log2 fold change (LFC) is also visualized along with the main heatmap. Genes with top LFC (> 3.5 or < -2) are highlighted. (C) Volcano plot of gene expression with thresholds (dashed lines) set at adjusted P value < 0.005 and absolute LFC ≥ 0.99. Names of the top 35 differentially expressed genes are presented. (D) GO enrichment analysis results for all significant and down-regulated genes with terms related to erythrocyte, lipid, and oxidative stress. The dot size reflects the gene numbers, and colors represent the q-values of these terms. Triangle dots indicate no significance (FDR q-value > 0.05). (E) The GSEA results of the erythrocyte-related GO terms. (F, G) Gene interaction networks related to erythrocyte and lipid regulation. The dot colors indicate the fold changes of the gene expression (blue: negative and red: positive). Created in BioRender. Min, J. (2025) https://BioRender.com/5mvwxqc

### *Scd1* is significantly downregulated in *Rpl5^+/-^* erythroid progenitors

We also performed RNA-seq using FL erythroid progenitors from single E12.5 embryos, comparing mutant and WT mice. We separated mutant samples into 2 groups based on the number of erythroid cells recovered, under the hypothesis that mutants with significantly lower cellularity in the erythroid progenitor gate (M-low) were likely to succumb to erythroid failure compared to mutants with higher cell counts (M-high) (Figure 4A). M-low embryos had lower numbers of cells in both quadrants II and III. To investigate the potential pathways involved in driving the cellularity differences between mutant groups, we performed the DE analysis among four types of comparisons between (1) M-low + M-high and wildtype (M-all vs. WT); (2) M-high and WT; (3) M-low and WT; (4) M-low and M-high. We identified 390, 231, and 407 DE genes in the WT comparisons (Figure 4B), with no significantly dysregulated noncoding RNAs. They were enriched in the GO terms relating to erythrocyte development and oxidative stress (Figures S4, S5). Five genes, including *Scd1*, were identified from the M-high vs. M-low comparison (Figure 4C, D). Notably, *Scd1* consistently downregulated across all comparisons, including the prior RNA-seq experiment presented in Figure 3 (Figure 4E). The same down-regulation of SCD (human homolog) was also observed in the previously published human data^39^ (LFC < -3.9, adjusted p-values < 1e-5, Figure S6), further suggesting its potential involvement in DBAS.

**Figure 4.**
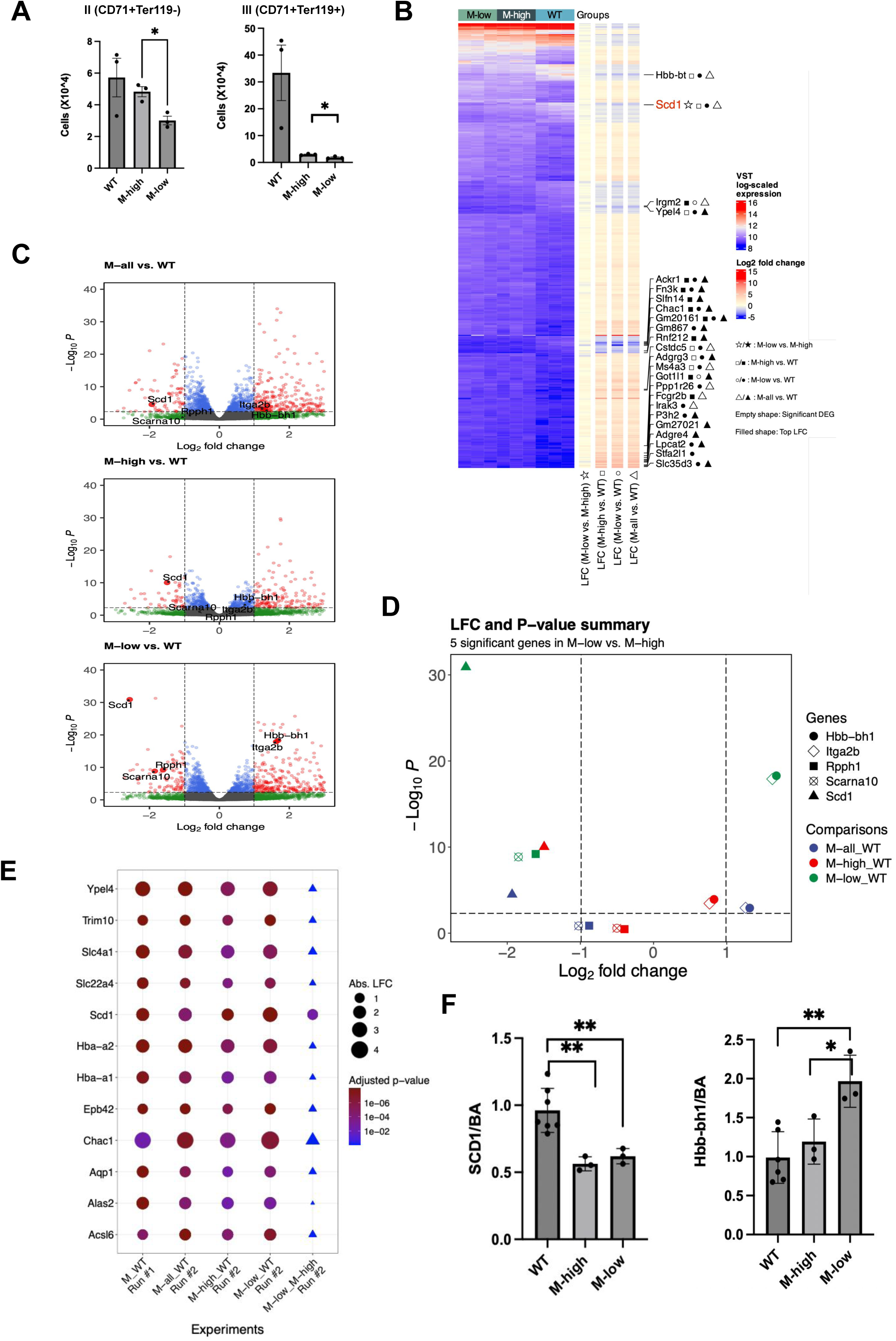
Mutant E12.5 FL with lower cellularity shows significant down-regulation of the lipid metabolism gene *Scd1.* We performed bulk RNA-seq using total RNA from sorted population II (CD71+ Ter119-) cells from E12.5 FL using single embryos (WT, n=3; *Rpl5^+/-^* n=6). Mutants were analyzed together and separately using differences in fetal liver erythroid cell counts obtained from sort (M-low, n=3; and M-high, n=3). (A) M-low embryos had significantly less erythroid progenitor cells in populations II (CD71+Ter119-) and III (CD71+Ter119+) compared with M-high. (B) The heatmap visualizes the variance stabilizing transformation (VST) log-scaled expressions for all the significant genes unioned from the four types of comparisons (M-high vs. WT, M-low vs. WT, M-all vs. WT, and M-low vs. M-high). Heatmap rows are automatically clustered when the columns are manually sorted. Corresponding LFC for the four comparisons are also visualized. Genes with top LFCs (≤ -2.5 or ≥ 5) from the four comparisons are labeled. The shapes after the gene names indicate in which comparisons they show significant differences and whether they have top LFCs. *Scd1* is highlighted as it is the only gene that shows significance across all four comparisons. (C) Volcano and (D) summary plot showing the comparisons between mutants and wild-types, highlighting the positions of the five significant genes in the M-low vs. M-high comparison. (E) Summary of common genes involved in the significant GO terms across all comparisons of the two RNA-seq experiments. Triangles indicate no significance (FDR q-value > 0.05). (F) Validation of significant genes in population II by RT-qPCR.

We validated dysregulated genes by RTq-PCR and demonstrated *Scd1* downregulation in E12.5 FL mutant erythroid progenitors (Figure 4F). *Hbb-bh1* was also significantly elevated in M-low but not M-high, which indicates that these two mutant groups have distinct erythroid features. SCD1 is known to catalyze the conversion of saturated to monounsaturated fatty acids^40^. However, to our knowledge, the role of SCD1 and lipid metabolism in erythropoiesis and DBAS has not been previously described.

### *Rpl5^Skax23-Jus/+^* adult mice have delayed erythroid recovery after stress

Our prior work in *Rpl5^Skax23-Jus/+^* mice demonstrated that postnatal macrocytic anemia leads to early mortality after birth in some mice, but surviving adult/aged mice exhibited no anemia or bone marrow failure. We hypothesized that the lack of anemia in adult animals might be partly due to the lack of hematological challenges (i.e., absence of infections/inflammation and/or environmental/dietary toxins) in normal mouse husbandry conditions, as seen in Fanconi mouse models, which do not develop bone marrow failure in the absence of stress^41,42^. To test this hypothesis, we first induced inflammatory stress with Poly(I:C), a synthetic double-stranded RNA that simulates a viral infection (Figure S7A). We found that Poly(I:C) induced significant anemia in both WT and *Rpl5^+/-^* mice by day 7. However, WT mice started to recover after day 7 despite being administered additional doses of Poly(I:C), while *Rpl5^+/-^* mice slowly recovered to baseline by day 28 (Figure S7B-E). We next tested a standard hematological stress condition in adult mice by injection of phenylhydrazine, which causes hemolysis (supplemental methods, Figure 5A). After phenylhydrazine administration, mutant mice showed an enhanced nadir and delayed recovery of RBC counts after the 1st and 2nd treatments, indicating that erythropoiesis in adult mutant mice was more severely affected by this hematological challenge (Figure 5B-C, Figure S8, S10). In this experiment, 3 out of 4 male *Rpl5^+/-^* mice treated with phenylhydrazine died, whereas there were no deaths in other groups (Figure S9).

**Figure 5.**
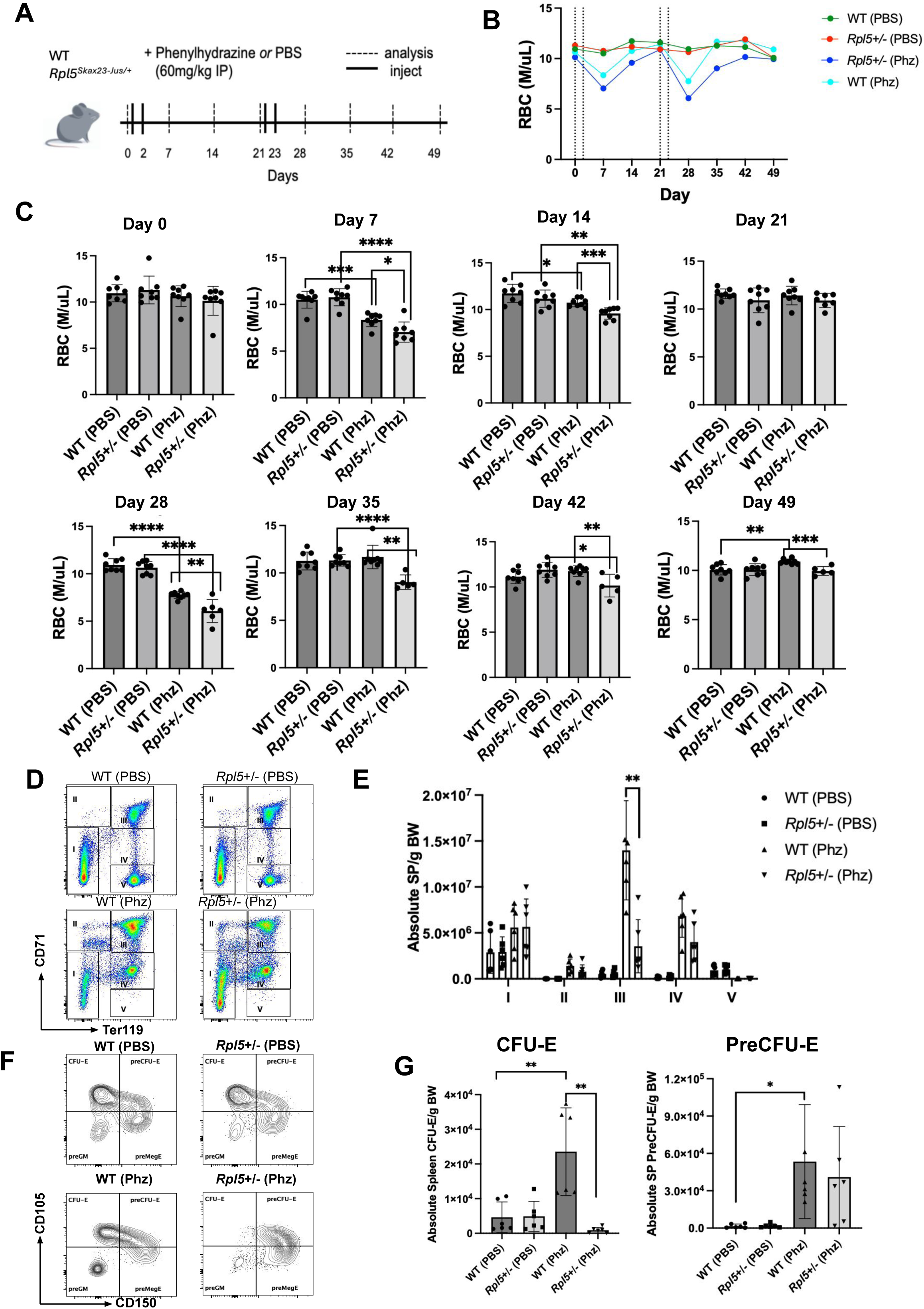
*Rpl5^Skax23-Jus/+^*mice show delayed erythroid recovery after phenylhydrazine treatment. (A) Adult (1.5-3.5 month) mice were injected with phenylhydrazine (Phz) (60µg/g IP) or PBS on days 0, 2, 21, and 23, and weekly blood count obtained until recovery (n=8 for each group:, 50/50 M:F ratio). (B) *Rpl5^+/-^* Phz-treated mice had a more significant macrocytic anemia with lower RBC after both Phz treatments compared with WT. WT and *Rpl5^+/-^* both showed recovery by days 21 and 49, however *Rpl5^+/-^* mice experienced slower recovery of RBC counts after Phz treatment and death occurred in 3 mutant mice (C) Graph of the RBC counts at each day of analysis in panel (B). (D) Analysis of erythropoiesis in the spleen using CD71 and Ter119 on day 4 with quantification in (E) (n=6 for each group:, 50/50 M:F ratio). (F) Analysis of CFU-E and PreCFU-E in the spleen on day 4 with quantification in (G).

In mice, stress erythropoiesis occurs predominantly in the spleen, which rapidly induces erythroid output in response to anemic stress^43^. We initially performed an analysis of peripheral blood counts days 3-5 post-phenylhydrazine administration and found that the RBC nadir occurred on day 4 (Figure S10). When we examined the spleen erythroblasts at day 4, CD71^+^Ter119^+^ cells in the spleen were reduced significantly in mutant mice (Figure 5D). We represented this data using total cells, absolute cells/g BW, and % live cells (Figures 5E, S10A) and showed a statistically significant decline in this progenitor population in all three analyses. There was an increase in the % live CD71^-^ Ter119^-^ population (population I) in *Rpl5^+/-^* mice but this was not statistically significant with absolute cell numbers. Bone marrow erythroid progenitors showed no change after phenylhydrazine treatment (Figure S11B). Next, we quantified early and late erythroid progenitors in the spleen after phenylhydrazine treatment (Figure S12A). In wildtype mice, the number of preMegE, preCFU-E, and CFU-E cells were elevated as early as four days after initial phenylhydrazine treatment (Figure 5F-G, Figure S12B-E), demonstrating efficient regeneration response. While the preMegE and preCFU-E numbers were not significantly impaired in *Rpl5^+/-^* adult spleen compared to WT spleens post-phenylhydrazine administration (Figure 5F-G, Figure S12C-D), the CFU-E population was profoundly diminished in mutant mice (Figure 5F-G, Figure S12B). This is the likely cause of delayed production of CD71^+^Ter119^+^ stress erythroid progenitors and delayed mature RBC production in the circulation after Poly(I:C) or phenylhydrazine treatment in mutant mice.

### An SCD1 inhibitor improves erythropoiesis in adult mice after anemic stress

We next addressed whether the aberrant pathways described in E12.5 FL mutant erythroid progenitors also played a role in adult stress erythropoiesis in our model. The specific question was whether *Scd1* downregulation drives the failure of erythropoiesis or if *Scd1* downregulation is a compensatory mechanism to augment erythropoiesis. We pretreated mice with an available SCD1 inhibitor (SCD1-i) 5m/kg (supplemental methods) or vehicle control (DMSO) for 14 days and then administered phenylhydrazine (Figure 6A). The mice showed some weight loss but no change in peripheral blood counts with SCD1-i (Figure S13A-B). There was a slight increase in bone marrow cellularity in WT mice treated with SCD1-i, but spleen cellularity was similar in all groups (Figure S13C). Mutant mice treated with vehicle control showed a significant decrease in Hgb and RBC compared with WT whereas SCD1-i-treated mice had no or significantly less differences in Hgb and RBC, respectively (Figure 6B). However, we terminated analysis after day 7 due to significant ocular toxicity from the drug.

**Figure 6.**
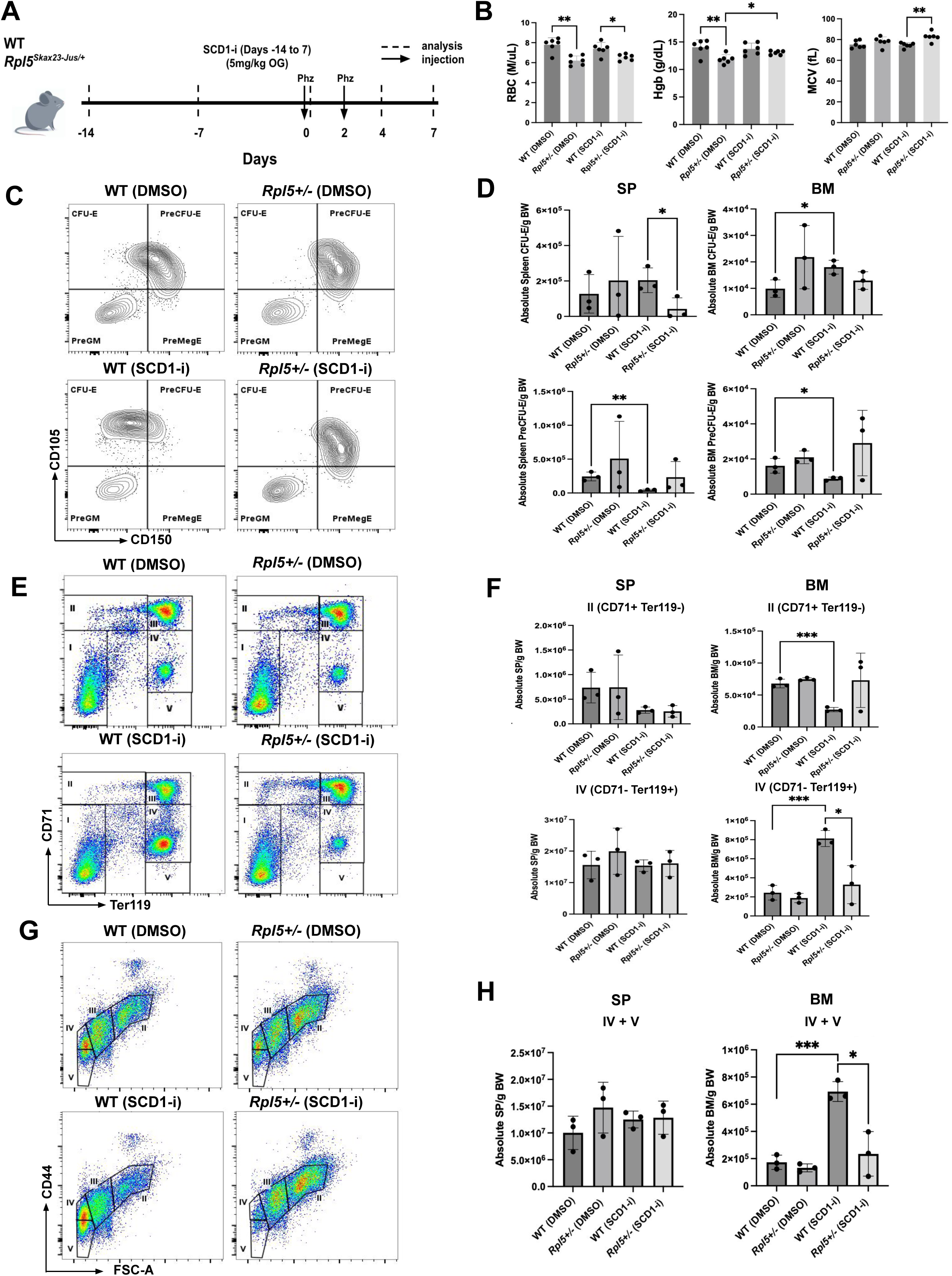
Treatment with SCD1 inhibitor improves erythropoiesis. (A) We pretreated adult mice (female, n=6 per group, age 2-4 months) with SCD1 inhibitor (CAY10566) or DMSO for two weeks (days -14-0) and then gave the mice phenylhydrazine on days 0 and 2. (B) Analysis of blood counts at day 7 showed a slight improvement in RBC counts in mutant mice after SCD1 treatment. Analysis of HSPC subsets by flow cytometry at day 4 (C, D) showed a significant increase in CFU-E and a decrease in pre-CFU-E in WT mice. (E, F) Analysis of erythroid precursors at day 4 by CD71 and Ter119 showed a decrease in earlier precursors (II) and an increase in late precursors (IV) in WT mice. (G, H) Further quantification of terminal erythropoiesis (Ter119+ gate) using CD44 vs FSC shows significant improvement in erythropoiesis in WT animals after SCD1-i treatment.

In order to explore the effect of SCD1 inhibition on erythropoiesis, we analyzed HSPC progenitors by flow cytometry. WT mice treated with SCD1-i showed a significant increase in CFU-E number in the bone marrow and decreased preCFU-E in both the bone marrow and spleen at day 4, compared to WT mice treated with DMSO (Figure 6C-D). There was no corresponding change in *Rpl5^+/-^* mice. Similarly, when we analyzed later erythroid progenitors, we saw improvement in erythropoiesis in WT mice treated with SCD1-i with a significant decrease in early CD71^+^Ter119^-^ (II) progenitors and an increase in late CD71^-^Ter119^+^ (IV) progenitors in bone marrow (Figure 6E-F). We did not observe a corresponding change in spleen erythroid progenitors. Analysis of terminal erythropoiesis (CD44 vs. FSC) showed a significant increase in mature bone marrow erythroid cells (IV + V) and an insignificant upward trend in *Rpl5^+/-^* mice (Figure 6G-H).

We next tested whether the lack of erythroid progenitor effect after SCD1 inhibition in mutant mice was due to drug toxicity or due to a lag in drug effect in mutant animals. Analysis at day 7 (Figure 7A) now showed a significant increase in spleen cellularity with the 1.25mg/kg SCD1-i dose in mutant mice with increased CFU-E expansion (Figure 7B), which was significant by CFU-E % (Figure 7C) with a trend towards significance in absolute spleen CFU-E counts/BW (Figure 7D). There was no significant effect in spleen CFU-E formation in mutant mice using 2.5mg/kg or 5mg/kg doses. Further analysis of erythropoiesis also showed a significant increase in CD71+Ter119+ cells in mutant mice after the 1.25mg/kg dose of SCD1-i (Figure 7E and F). This indicates that erythropoiesis in *Rpl5* haploinsufficient mice is responsive to SCD1 inhibition but at lower doses and with a longer response time than required for WT mice.

**Figure 7.**
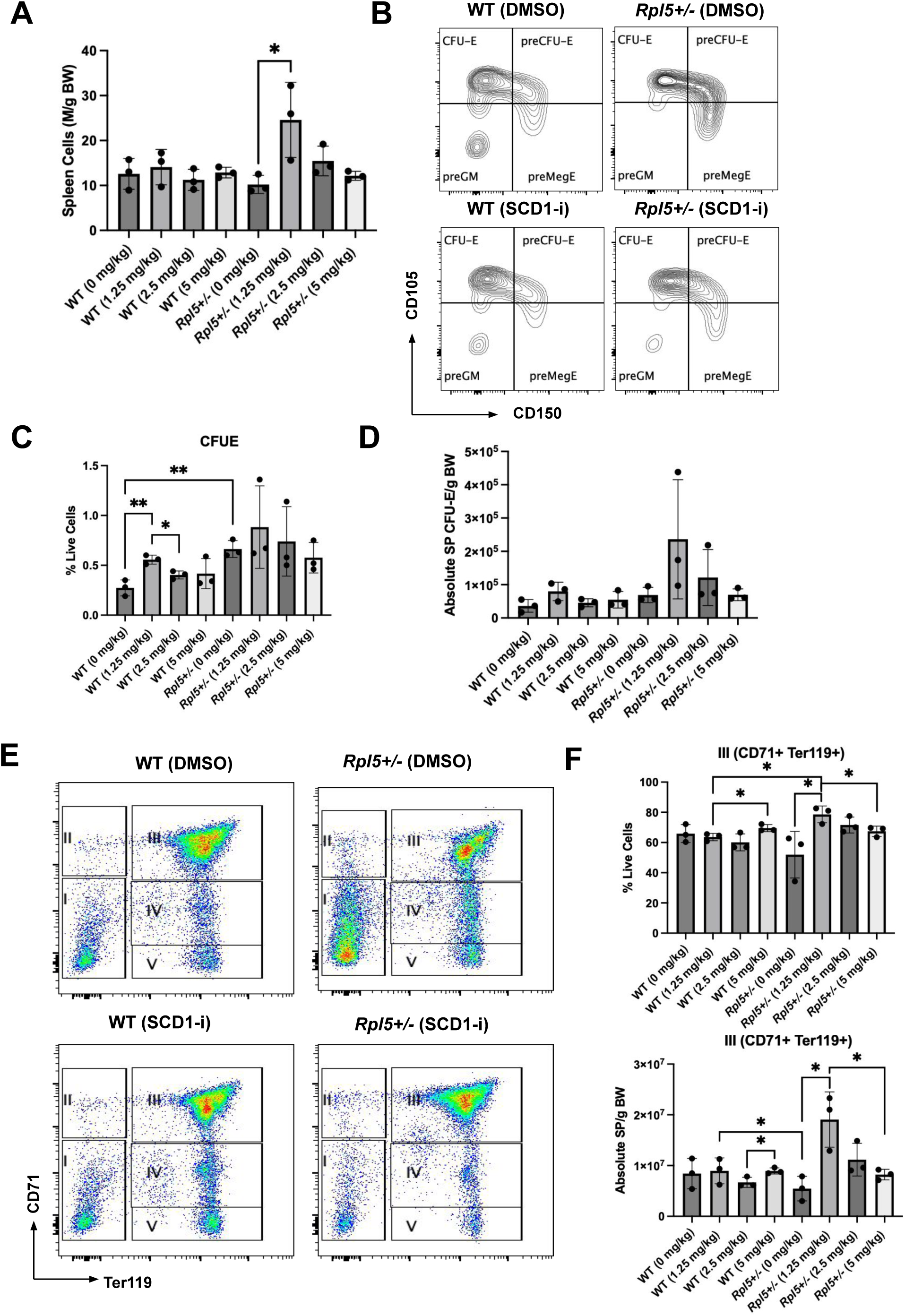
SCD1 plays a role in stress erythropoiesis in *Rpl5^Skax23-Jus/+^* mice. Adult mice were pretreated with SCD1 inhibitor (1.25mg/kg, 2.5mg/kg or 5mg/kg) for 2 weeks and then given phenylhydrazine at days 0 and day 2. Spleen cellularity was obtained at day 7 (A) and quantification of spleen HPSC progenitors obtained by flow cytometry (B-D), which demonstrated a trend towards increased CFU-E and pre-CFU-E in mutant animals with SCD1 inhibitor at the 1.25mg/kg dose. (E, F) Further analysis demonstrated improvement of erythropoiesis in mutant animals with reduction in CD71-Ter119-progenitors (I) and increase in CD71+ Ter119+ erythroid cells (III).

## Discussion

This work has further characterized *Rpl5^Skax23-Jus/+^* mice, which we initially described as an accurate model of the variable erythroid phenotype in DBA syndrome. Here, we describe additional salient features of this mouse that align with the phenotype seen in the human disorder. DBAS patients with *RPL5 and RPL11* mutations have been noted to have a higher incidence of congenital malformations including craniofacial defects^2^. We did not initially detect a postnatal craniofacial defect in the mutant mice, but in subsequent analysis of more embryos, we identified the presence of a subset of mice with abnormal palate development. The craniofacial defects did not always occur in the VSD setting, and surviving adult mice had no evidence of either defect. The data indicate that these morphological defects are markers of a severe disease phenotype and suggest that *Rpl5* haploinsufficiency may affect neural crest cells and/or downstream progenitors in a stochastic manner^44,45^. We further found an elevation of proliferating cells and a reduction in the number of apoptotic cells in *Rpl5^+/-^* mice compared to controls. One of the functions of RPL5 is to activate p53 functions by suppressing MDM2, an inhibitor of p53. In prior work, we demonstrated that cells haploinsufficient for *Rpl5* do not show aberrant basal p53 activation and had a p53-independent cell cycle defect, consistent with a nucleolar stress model in which RPL5 plays a crucial role in the 5S RNP complex^46,47^. Thus, the elevation of proliferation and reduction of cell death in *Rpl5* haploinsufficient mice are expected outcomes due to the suppression of normal p53 regulation. Retinoic acid-induced cleft palate, a major known causative reason for cleft palate, also showed an elevation of proliferating cells^48^. These data suggest that embryos with *Rpl5* haploinsufficiency may be at risk for development of craniofacial anomalies such as cleft palate and lip through abnormal proliferation and cell death caused by suppression of normal p53 functions during embryogenesis. Further work is needed to determine the signaling pathways that drive the variable development defects in *RPL5* haploinsufficient DBAS as well as in other genotypes that exhibit p53 activation.

Analysis of DBAS cohorts has revealed a wide spectrum of genetic variants leading to ribosome biogenesis defects from canonical loss of function, missense or splice site mutations in ribosomal proteins or in ribosome-associated proteins such as *HEATR3* and *TSR2*^2^. About 20% of patients do not have a known ribosomal protein variant. Some of these patients have been subsequently found to have non-canonical splice or deep intronic mutations while others have non-ribosome variants such as in *GATA1* or gain-of-function *TP53* mutations, which share elements of the defective pathways seen in RP-haploinsufficient DBAS. The mutation seen in *Rpl5^Skax23-Jus/+^* mice is at a common splice site, which should be identified by exome sequencing. However deep intronic mutations can affect gene function and may be missed by typical exome sequencing. Whole genome analysis and/or functional analysis of ribosomal levels may be warranted in those cases but have limited availability to clinicians^49,50^. Others have proposed use of minigene assays to further characterize variants of unknown significance, which occur frequently in clinical genetic testing of rare diseases^51^. Here we demonstrate that the pathogenesis of this variant can also be detected by intron retention bioinformatic analysis of RNA-seq data.

Several mechanisms have been proposed to explain the erythropoiesis defect in DBAS, which are not mutually exclusive^52^. However, these mechanisms do not offer an explanation for the spontaneous occurrence of treatment-independent periods or a lack of anemia in some patients. In our model, this resolution of anemia occurred quite rapidly within the same litter during the period from E12.5 to weaning, which argues against major acquired genetic/clonal processes and environmental factors but may indicate a role of metabolic and/or epigenetic factors^53^. As some DBAS patients have anemia recurrence during stressful periods such as pregnancy and viral infections^7^, we sought to replicate this disease aspect in the current animal model. We demonstrated that adult mutant mice without anemia had re-induction of anemia with additional stress and exhibited a delay in erythroid recovery, modeled here with both poly(I:C) and phenylhydrazine. This argues that DBAS may not be a *de novo* anemia syndrome but perhaps a syndrome characterized by defective or delayed recovery from erythropoietic stress as demonstrated by the findings here. Analysis of the pathways that support anemia resolution and/or that augment stress erythropoiesis are therefore critical to further understand this disease aspect, which may aid in rational therapeutic strategies.

Advances in “omics” have recently facilitated our understanding of the role of metabolic regulation in erythropoiesis. Mitochondria, amino acid, iron/heme, glycolysis and lipid metabolism are all involved in the stepwise process of erythroid lineage commitment, differentiation, and maturation^54^. Among these, the regulation of lipid metabolism during erythropoiesis has not been well-studied. Metabolomic analysis of cells undergoing terminal erythropoiesis revealed the key role of PHOSPHO1 in regulating phosphocholine metabolism, ATP production, and amino acid supply during erythropoiesis^55^, expanding the knowledge of the link between lipid composition and erythropoiesis. Prior work in DBAS cohorts and models have described various metabolic abnormalities. For example, metabolic fingerprints of dried blood spot samples identified increased inosine in DBAS patients compared with controls and elevated alpha-tocopherol levels in treatment-independent DBAS patients^56^; metabolic profiles revealed increased amino acid metabolism or nucleotide pool depletion in different DBAS individuals with distinct *RPL9* variants^57^. However, further validations of these metabolic disturbances were lacking in those studies. Here, we identified the downregulation of *Scd1* through unbiased analysis and verified the role of lipid metabolism in promoting erythropoiesis with a known SCD1 inhibitor. These findings shed light on the association between lipid metabolism and erythropoiesis.

Stearoyl-CoA desaturase 1 (SCD1) is a critical enzyme that desaturates saturated fatty acids to monounsaturated fatty acids (MUFAs), therefore maintaining homeostasis of lipid metabolism^58^. Aberrant SCD1 expression can cause lipid accumulation, promoting metabolism-related diseases such as obesity and diabetes. In addition, SCD1 plays a role in oncogenesis and SCD1 inhibition is currently being examined in clinical trials^40^. Despite a large body of prior work in this area, the role of SCD1 in erythropoiesis was not previously described. Here, we found significant downregulation of *Scd1* in M-low embryos, which we propose are the mutant group likely to succumb to failure of erythropoiesis. A key question here was whether *Scd1* downregulation was the cause or consequence of these erythropoietic defects. By examining the effect of SCD1 inhibitor treatment in the PHZ-induced anemic mouse model, we demonstrated that SCD1 inhibition promoted erythroid maturation, suggesting that downregulation of *Scd1* is likely a compensatory mechanism to augment erythropoiesis. Interestingly, this effect required a lower dose and a longer response time in *Rpl5* haploinsufficient mice, possibly due to the already low expression level of *Scd1* in these mutants. Our future work will focus on understanding the specific role of lipid metabolism and SCD1 during normal and defective erythropoiesis in Diamond Blackfan anemia syndrome, which we expect will increase our fundamental understanding of erythropoiesis and guide therapeutic approaches.

## Data availability

Raw sequencing data and the processed expression matrix in raw counts can be accessed via GSE297106.

## Acknowledgments

Research reported in this publication was supported by the National Institutes of Health grant P30CA046592 to the University of Michigan Rogel Cancer Center, National Institutes of Health grants K08 DK127013 (S.A.S), R01 HL148333 (R.K.), R01 HL157062 (R.K.), U2C DK129445 (R.K.) and American Heart Association Grant #923720/Lumeng/2021. This work was also supported by University of Michigan grants (S.A.S.): Janette Ferrantino Investigator, Loeb, and Holden Awards. We acknowledge support from the Bioinformatics Core of the University of Michigan Medical School’s Biomedical Research Core Facilities (RRID:SCR_019168).

## Authorship Contributions

S.A.S, J.D.E and R.K conceived the study and designed the experiments. Y.W, G.M, L.Y, X.L, S.H, J.C.M, A.A, R.S, V.N, H.U, J.L.Y, V.K, S.A.S performed experiments and analyzed the experimental data. K.D and Y.G performed bioinformatic analysis. C.D, B.C, J.K, C.L, Y.M, M.J, R.K, J.D.E contributed to results analysis. S.A.S, Y.W, K.D wrote the manuscript with assistance from all authors. All authors contributed to the evaluation, integration and discussion of the results.

