## Supplemental Figures for "A lipid metabolism defect is an underlying contributor to Diamond Blackfan anemia syndrome"

***Table S1***

| Antibody | Supplier | Catalog number |
|---|---|---|
| APC anti-HbF | Fisher Scientific | MHFH05 |
| FITC anti-humanCD71 | Biolegend | 334104 |
| APC anti-humanCD235a | BD Biosciences | 551336 |
| FITC anti-CD71 | Biolegend | 113806 |
| APC anti-Ter119 | Biolegend | 116212 |
| APC/Cy7 anti-CD44 | Biolegend | 103028 |
| Zombie Aqua | Biolegend | 423101 |
| BV421 anti-Sca1 | Biolegend | 108128 |
| APC anti-CD150 | Biolegend | 115910 |
| APC/Cy7 anti-CD117 | Biolegend | 105826 |
| Alexa Fluor700 anti-CD41 | Biolegend | 133926 |
| PE/Cy7 anti-CD105 | Biolegend | 120410 |
| BV605 anti-CD16/32 | BD Horizon | 563006 |
| FITC anti-CD48 | Biolegend | 103403 |
| Biotin anti-CD11b | Biolegend | 101204 |
| Biotin anti-Gr1 | Biolegend | 108404 |
| Biotin anti-B220 | Biolegend | 103204 |
| Biotin anti-CD3 | Biolegend | 100304 |
| Biotin anti-CD4 | Biolegend | 100404 |
| Biotin anti-CD5 | Biolegend | 100604 |
| Biotin anti-CD8 | Biolegend | 100704 |
| Biotin anti-TER119 | Biolegend | 116204 |
| PE-Cy5 anti-Streptavidin | Biolegend | 405205 |

***Table S2***

| **Oligonucleotide Name** | **Sequence (5'-3')** |
|---|---|
| Ms SCD1 F | GCAAGCTCTACACCTGCCTCTT |
| Ms SCD1 R | CGTGCCTTGTAAGTTCTGTGGC |
| Ms Rpl5 F | GCGCTACCTAATGGAGGAAGATG |
| Ms Rpl5 R | CTCTCGGATAGCAGCATGAGCT |
| Ms BA F | GAGGCATACAGGGACAGCAC |
| Ms BA R | CTAAGGCCAACCGTGAAAAG |
| Ms Hbb-bh1F | ATCATGGGAAACCCCCGGA |
| Ms Hbb-bh1F | GGGTGAATTCCTTGGCAAAATGAGT |

***Table S3***

| Sample# | Genotype | VSD | Cleft | Palate Fusion | Description |
|---|---|---|---|---|---|
| (3-1) | *Rpl5*$^{+/-}$ | No | No | Yes | |
| (3-2) | WT | No | No | Yes | |
| (3-3) | *Rpl5*$^{+/-}$ | No | No | Yes | |
| (3-4) | *Rpl5*$^{+/-}$ | No | No | Yes | |
| (3-5) | *Rpl5*$^{+/-}$ | No | No | Yes | |
| (3-6) | WT | No | No | Yes | |
| (3-7) | *Rpl5*$^{+/-}$ | Yes | No | Yes | |
| (6-1) | *Rpl5*$^{+/-}$ | No | Yes | No | 1 shelf unelevated |
| (6-2) | *Rpl5*$^{+/-}$ | No | No | Yes | |
| (6-3) | WT | No | No | Yes | |
| (6-4) | WT | No | No | Yes | |
| (6-5) | WT | No | No | Yes | |
| (6-6) | *Rpl5*$^{+/-}$ | Yes | Yes | No | elevated, no fusion |
| (6-7) | WT | No | No | Yes | |
| (10-1) | WT | No | No | Yes | |
| (10-2) | *Rpl5*$^{+/-}$ | Yes | Yes | No | both shelves unelevated |
| (10-3) | WT | No | No | Yes | |
| (32-1) | WT | No | No | Yes | |
| (32-2) | *Rpl5*$^{+/-}$ | No | No | Yes | |
| (32-3) | *Rpl5*$^{+/-}$ | No | Yes | No | 1 shelf unelevated |
| (32-4) | *Rpl5*$^{+/-}$ | No | No | Yes | |
| (32-5) | WT | No | No | Yes | |
| (32-6) | WT | No | No | Yes | |
| (32-7) | *Rpl5*$^{+/-}$ | No | No | Yes | |

#### *Figure S1*

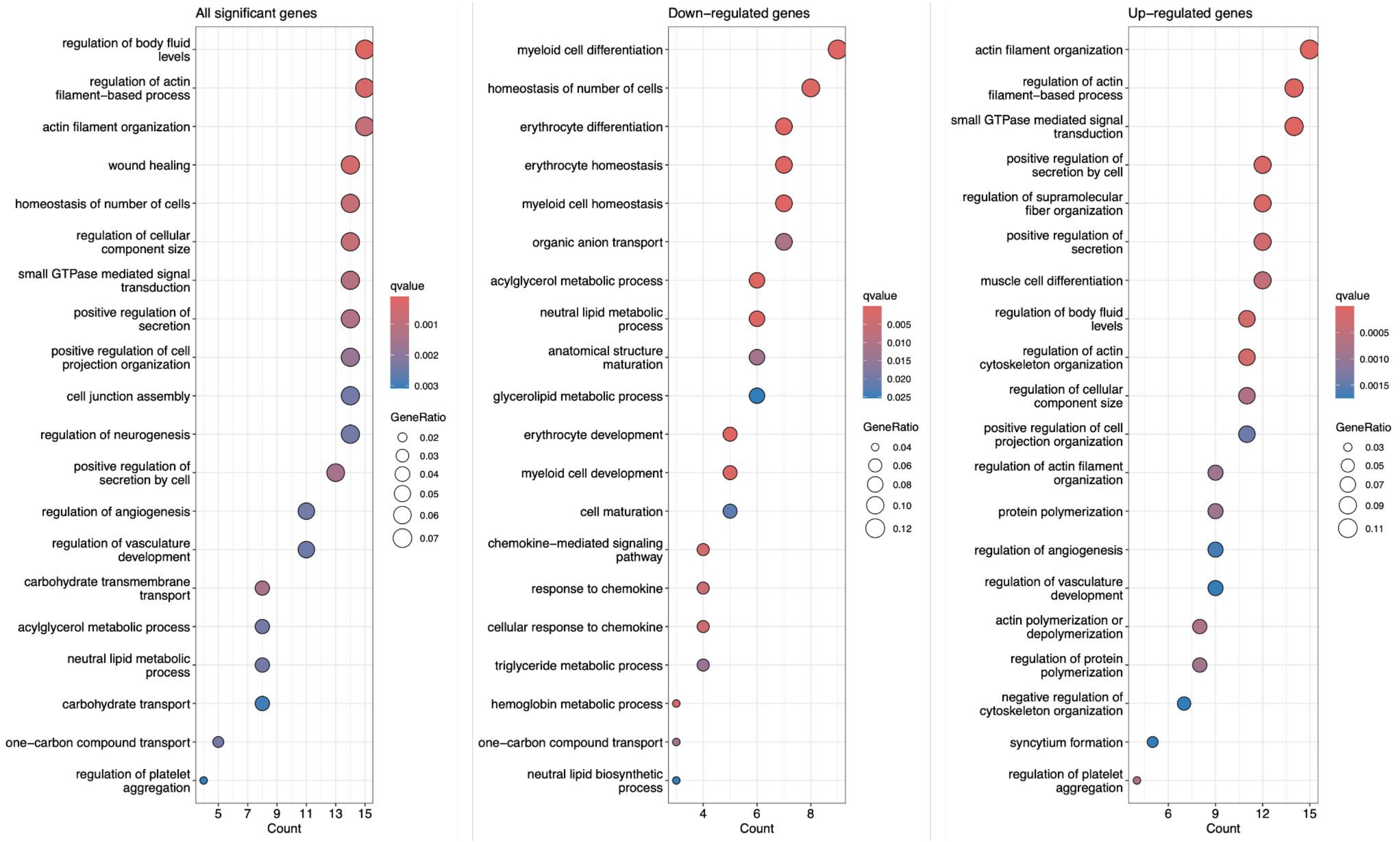
All significant genes
regulation of body fluid levels
regulation of actin filament−based process
actin filament organization
wound healing
homeostasis of number of cells
regulation of cellular component size
small GTPase mediated signal transduction
positive regulation of secretion
positive regulation of cell projection organization
cell junction assembly
regulation of neurogenesis
positive regulation of secretion by cell
regulation of angiogenesis
regulation of vasculature development
carbohydrate transmembrane transport
acylglycerol metabolic process
neutral lipid metabolic process
carbohydrate transport
one−carbon compound transport
regulation of platelet aggregation
5 7 9 11 13 15
Count
qvalue
0.001
0.002
0.003
GeneRatio
0.02
0.03
0.04
0.05
0.06
0.07
Down−regulated genes
myeloid cell differentiation
homeostasis of number of cells
erythrocyte differentiation
erythrocyte homeostasis
myeloid cell homeostasis
organic anion transport
acylglycerol metabolic process
neutral lipid metabolic process
anatomical structure maturation
glycerolipid metabolic process
erythrocyte development
myeloid cell development
cell maturation
chemokine−mediated signaling pathway
response to chemokine
cellular response to chemokine
triglyceride metabolic process
hemoglobin metabolic process
one−carbon compound transport
neutral lipid biosynthetic process
4 6 8
Count
qvalue
0.005
0.010
0.015
0.020
0.025
GeneRatio
0.04
0.06
0.08
0.10
0.12
Up−regulated genes
actin filament organization
regulation of actin filament−based process
small GTPase mediated signal transduction
positive regulation of secretion by cell
regulation of supramolecular fiber organization
positive regulation of secretion
muscle cell differentiation
regulation of body fluid levels
regulation of actin cytoskeleton organization
regulation of cellular component size
positive regulation of cell projection organization
regulation of actin filament organization
protein polymerization
regulation of angiogenesis
regulation of vasculature development
actin polymerization or depolymerization
regulation of protein polymerization
negative regulation of cytoskeleton organization
syncytium formation
regulation of platelet aggregation
6 9 12 15
Count
qvalue
0.0005
0.0010
0.0015
GeneRatio
0.03
0.05
0.07
0.09
0.11

*Figure S2*

**A**

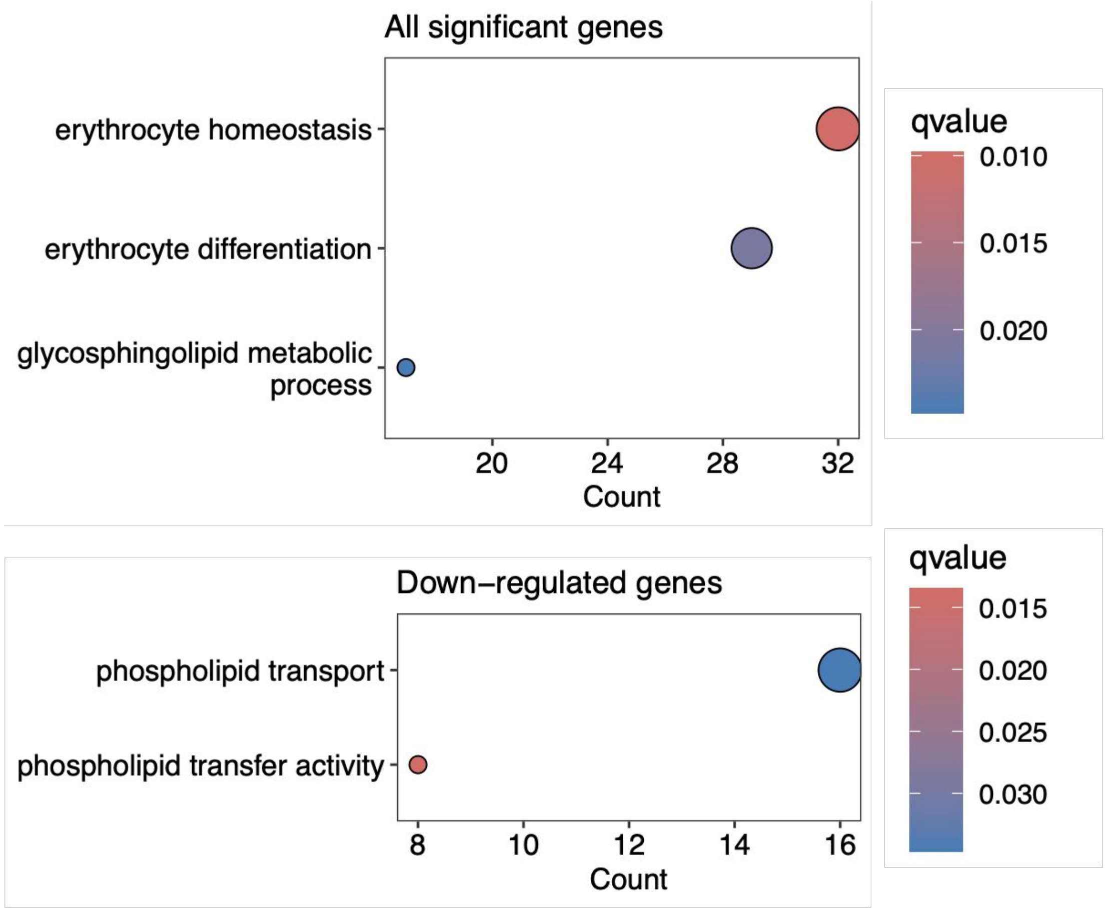
All significant genes
erythrocyte homeostasis
erythrocyte differentiation
glycosphingolipid metabolic process
20
24
28
32
Count
qvalue
0.010
0.015
0.020
Down−regulated genes
phospholipid transport
phospholipid transfer activity
8
10
12
14
16
Count
qvalue
0.015
0.020
0.025
0.030

**B**

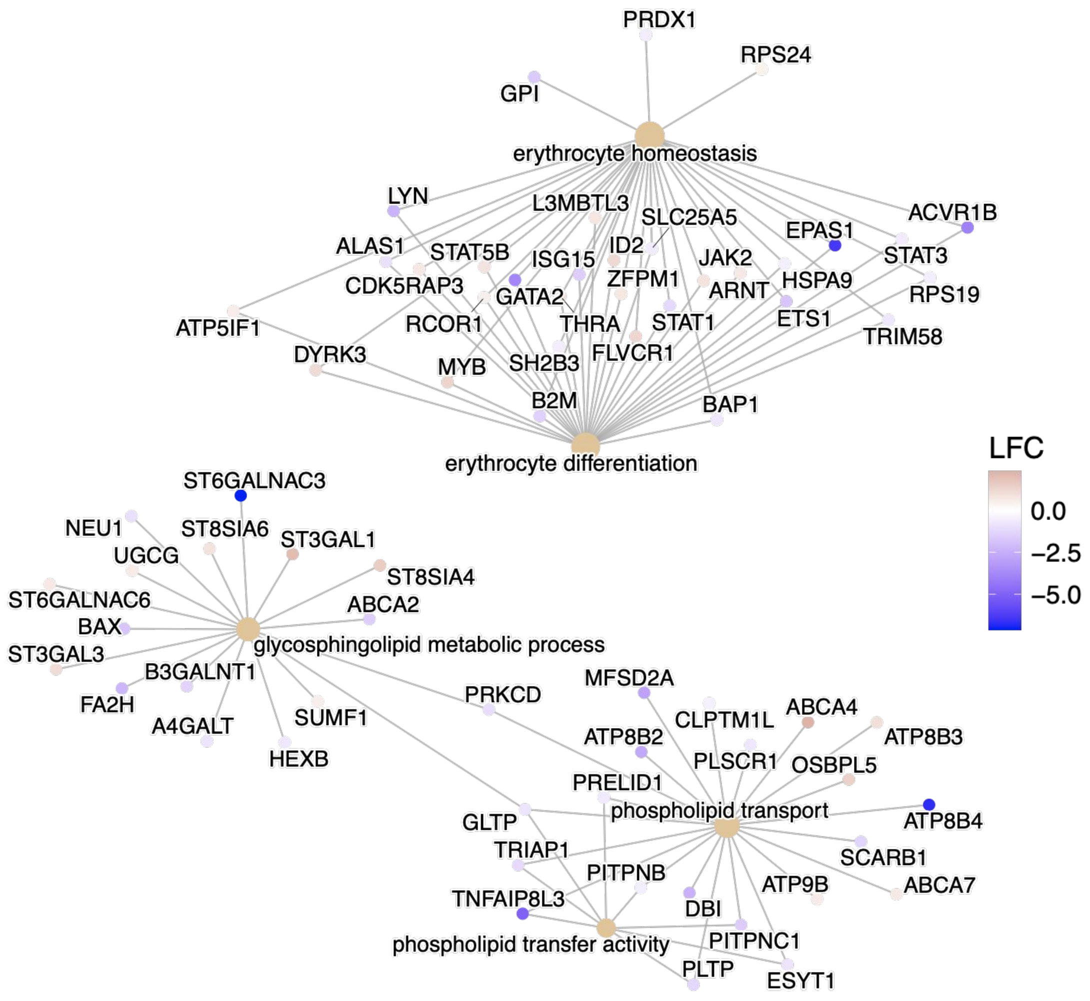
PRDX1
RPS24
GPI
erythrocyte homeostasis
LYN
L3MBTL3
SLC25A5
ACVR1B
EPAS1
ALAS1
STAT5B
ID2
ISG15
JAK2
STAT3
CDK5RAP3
ZFPM1
HSPA9
GATA2
ARNT
RPS19
ATP5IF1
RCOR1
THRA
STAT1
ETS1
TRIM58
DYRK3
MYB
SH2B3
FLVCR1
B2M
BAP1
erythrocyte differentiation
LFC
0.0
−2.5
−5.0
ST6GALNAC3
NEU1
ST8SIA6
ST3GAL1
UGCG
ST8SIA4
ST6GALNAC6
ABCA2
BAX
ST3GAL3
glycosphingolipid metabolic process
B3GALNT1
FA2H
A4GALT
SUMF1
HEXB
PRKCD
MFSD2A
CLPTM1L
ABCA4
ATP8B2
ATP8B3
PLSCR1
OSBPL5
PRELID1
phospholipid transport
GLTP
ATP8B4
TRIAP1
SCARB1
PITPNB
ATP9B
ABCA7
TNFAIP8L3
DBI
phospholipid transfer activity
PITPNC1
PLTP
ESYT1

*Figure S3*

**A**

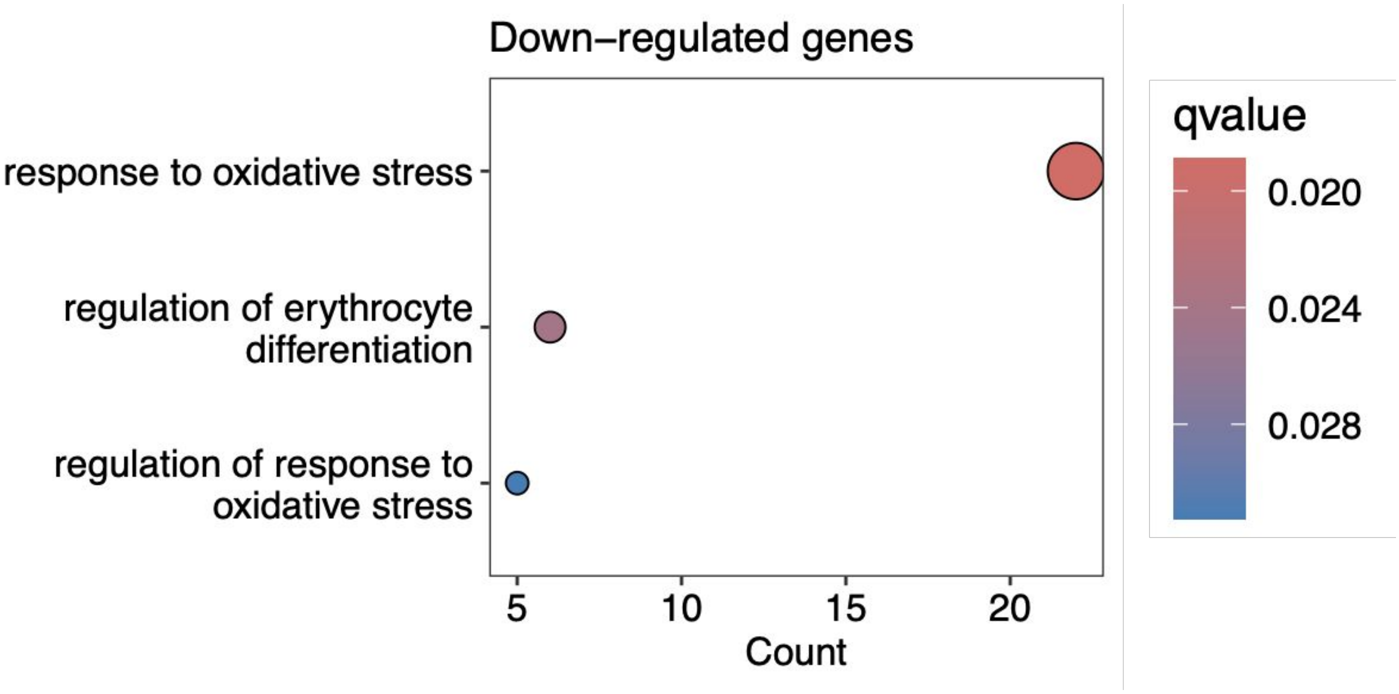

Down–regulated genes
response to oxidative stress
regulation of erythrocyte differentiation
regulation of response to oxidative stress
5
10
15
20
Count
qvalue
0.020
0.024
0.028

**B**

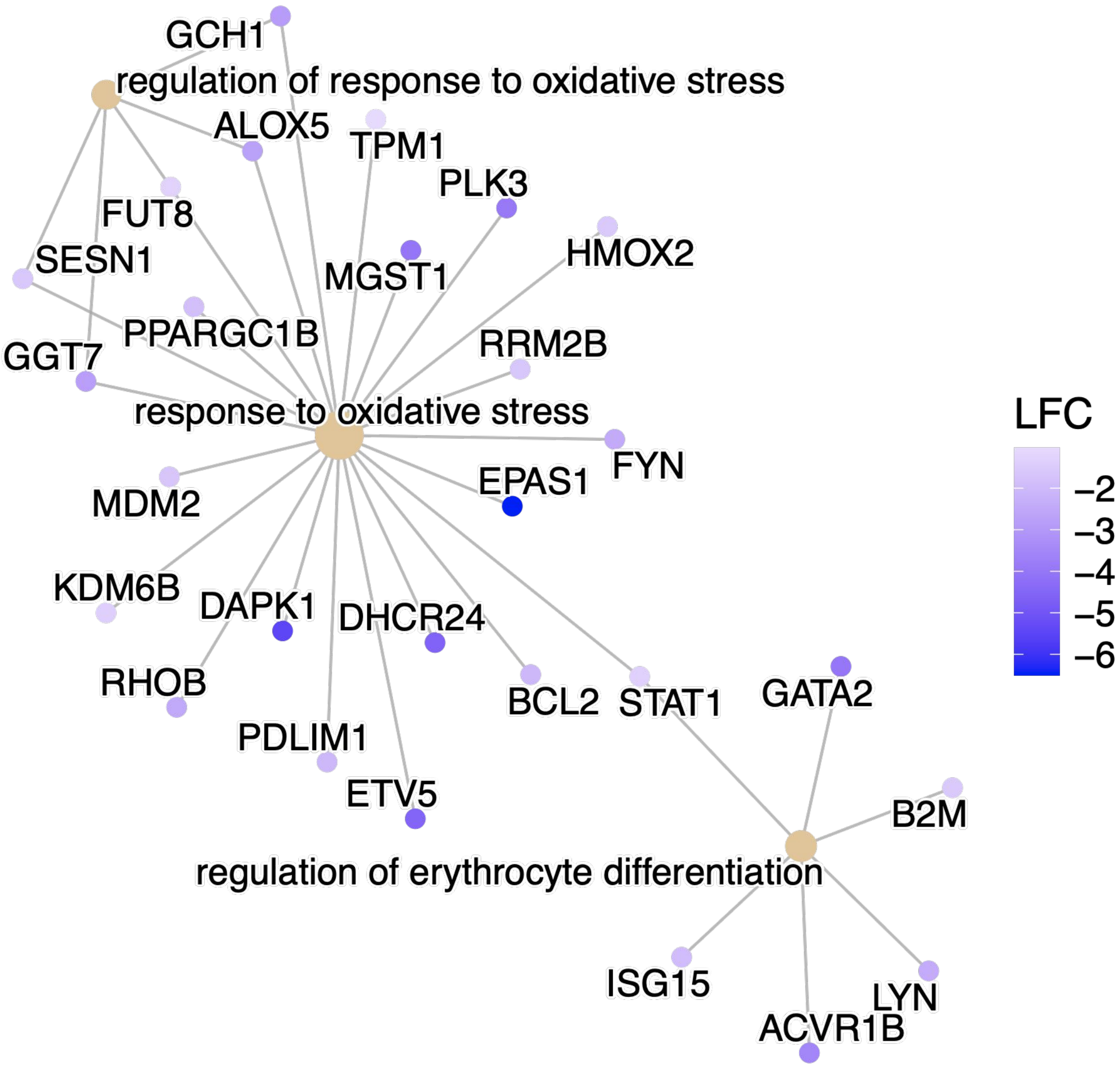

GCH1
regulation of response to oxidative stress
ALOX5
TPM1
PLK3
FUT8
SESN1
MGST1
HMOX2
PPARGC1B
GGT7
RRM2B
response to oxidative stress
FYN
EPAS1
MDM2
KDM6B
DAPK1
DHCR24
RHOB
BCL2
STAT1
GATA2
PDLIM1
ETV5
B2M
regulation of erythrocyte differentiation
ISG15
LYN
ACVR1B
LFC
−2
−3
−4
−5
−6

***Figure S4***

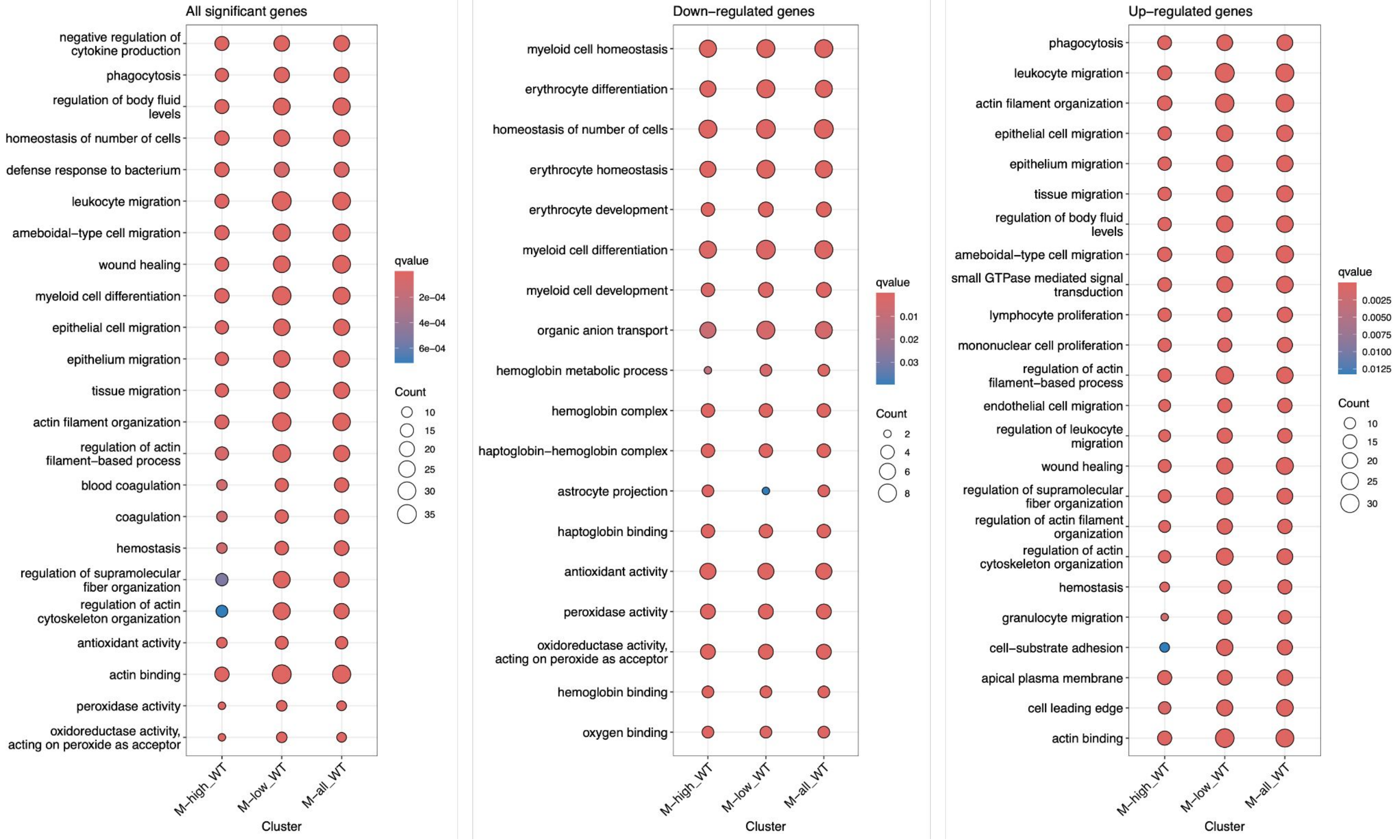
All significant genes
negative regulation of cytokine production
phagocytosis
regulation of body fluid levels
homeostasis of number of cells
defense response to bacterium
leukocyte migration
ameboidal−type cell migration
wound healing
myeloid cell differentiation
epithelial cell migration
epithelium migration
tissue migration
actin filament organization
regulation of actin filament−based process
blood coagulation
coagulation
hemostasis
regulation of supramolecular fiber organization
regulation of actin cytoskeleton organization
antioxidant activity
actin binding
peroxidase activity
oxidoreductase activity, acting on peroxide as acceptor
M−high_WT
M−low_WT
M−all_WT
Cluster
qvalue
2e−04
4e−04
6e−04
Count
10
15
20
25
30
35
Down−regulated genes
myeloid cell homeostasis
erythrocyte differentiation
homeostasis of number of cells
erythrocyte homeostasis
erythrocyte development
myeloid cell differentiation
myeloid cell development
organic anion transport
hemoglobin metabolic process
hemoglobin complex
haptoglobin−hemoglobin complex
astrocyte projection
haptoglobin binding
antioxidant activity
peroxidase activity
oxidoreductase activity, acting on peroxide as acceptor
hemoglobin binding
oxygen binding
M−high_WT
M−low_WT
M−all_WT
Cluster
qvalue
0.01
0.02
0.03
Count
2
4
6
8
Up−regulated genes
phagocytosis
leukocyte migration
actin filament organization
epithelial cell migration
epithelium migration
tissue migration
regulation of body fluid levels
ameboidal−type cell migration
small GTPase mediated signal transduction
lymphocyte proliferation
mononuclear cell proliferation
regulation of actin filament−based process
endothelial cell migration
regulation of leukocyte migration
wound healing
regulation of supramolecular fiber organization
regulation of actin filament organization
regulation of actin cytoskeleton organization
hemostasis
granulocyte migration
cell−substrate adhesion
apical plasma membrane
cell leading edge
actin binding
M−high_WT
M−low_WT
M−all_WT
Cluster
qvalue
0.0025
0.0050
0.0075
0.0100
0.0125
Count
10
15
20
25
30

*Figure S5*

**A**

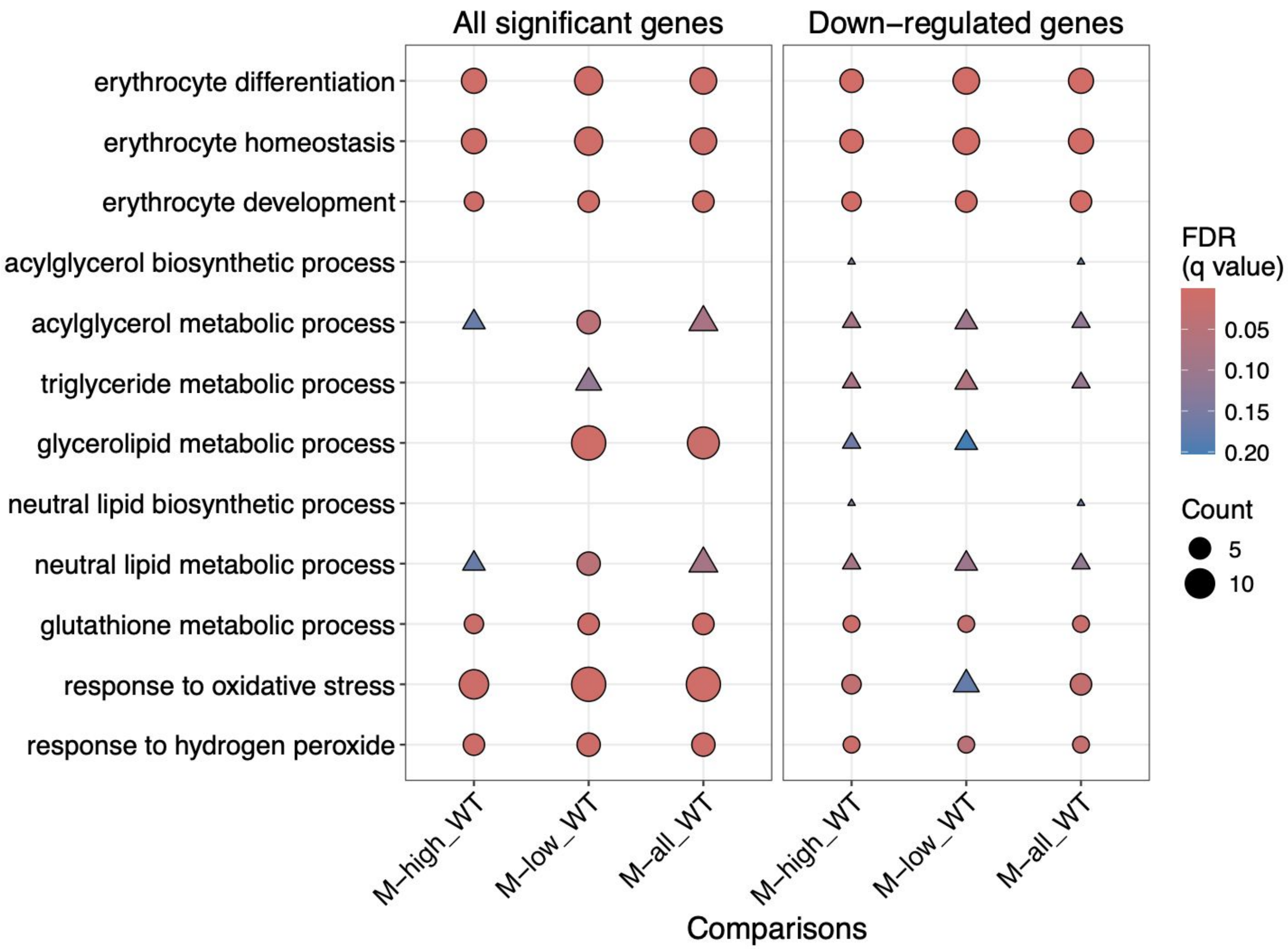
All significant genes
Down–regulated genes
erythrocyte differentiation
erythrocyte homeostasis
erythrocyte development
acylglycerol biosynthetic process
acylglycerol metabolic process
triglyceride metabolic process
glycerolipid metabolic process
neutral lipid biosynthetic process
neutral lipid metabolic process
glutathione metabolic process
response to oxidative stress
response to hydrogen peroxide
FDR
(q value)
0.05
0.10
0.15
0.20
Count
5
10
M–high_WT
M–low_WT
M–all_WT
M–high_WT
M–low_WT
M–all_WT
Comparisons

**B**

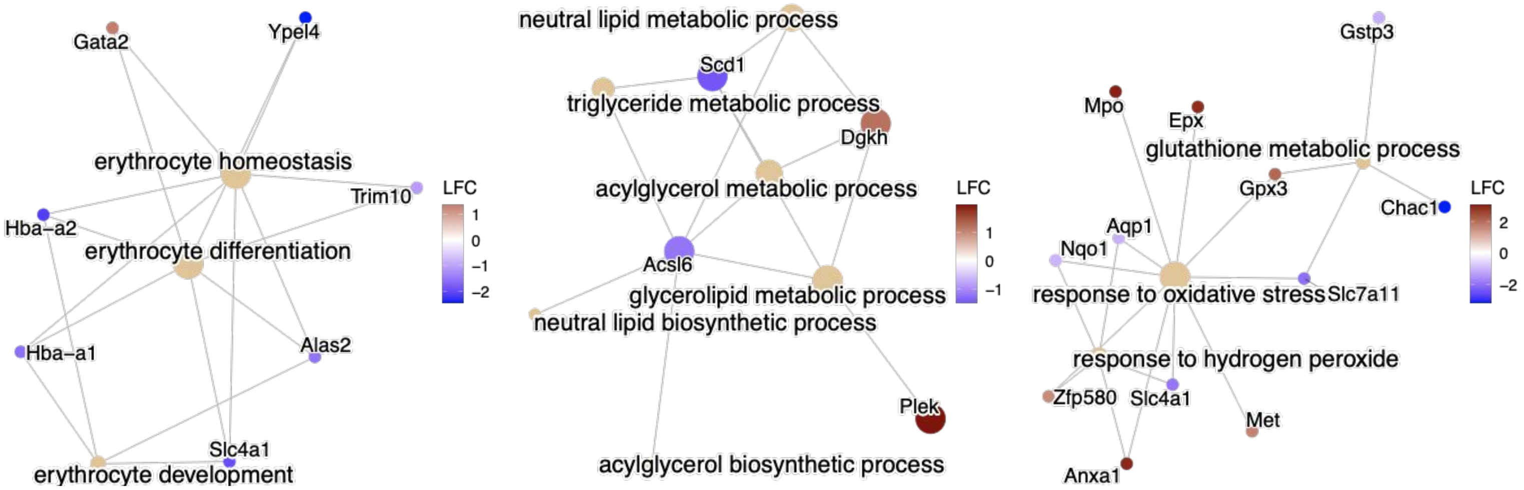
Gata2
Ypel4
erythrocyte homeostasis
Trim10
Hba–a2
erythrocyte differentiation
Hba–a1
Alas2
Slc4a1
erythrocyte development
LFC
1
0
−1
−2
neutral lipid metabolic process
Scd1
triglyceride metabolic process
Dgkh
acylglycerol metabolic process
Acsl6
glycerolipid metabolic process
neutral lipid biosynthetic process
Plek
acylglycerol biosynthetic process
LFC
1
0
−1
Gstp3
Mpo
Epx
glutathione metabolic process
Gpx3
Chac1
Aqp1
Nqo1
response to oxidative stress
Slc7a11
response to hydrogen peroxide
Zfp580
Slc4a1
Met
Anxa1
LFC
2
0
−2

*Figure S6*

A

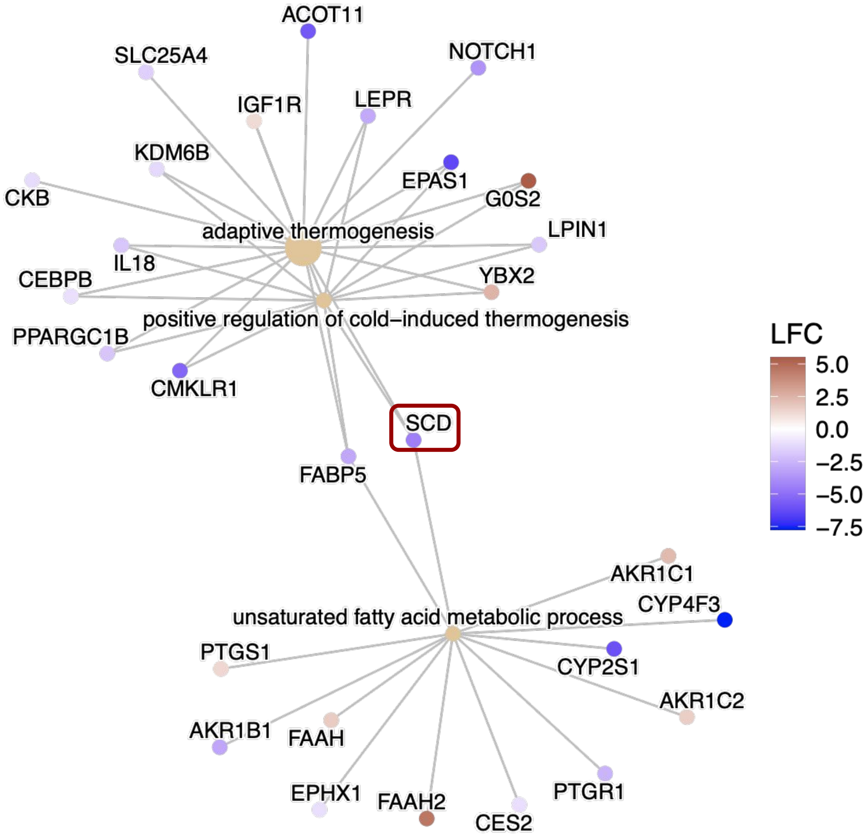
ACOT11
SLC25A4
NOTCH1
IGF1R
LEPR
KDM6B
EPAS1
CKB
G0S2
adaptive thermogenesis
LPIN1
IL18
CEBPB
YBX2
positive regulation of cold−induced thermogenesis
PPARGC1B
CMKLR1
SCD
FABP5
LFC
5.0
2.5
0.0
−2.5
−5.0
−7.5
AKR1C1
CYP4F3
unsaturated fatty acid metabolic process
PTGS1
CYP2S1
AKR1C2
AKR1B1
FAAH
EPHX1
FAAH2
CES2
PTGR1

B

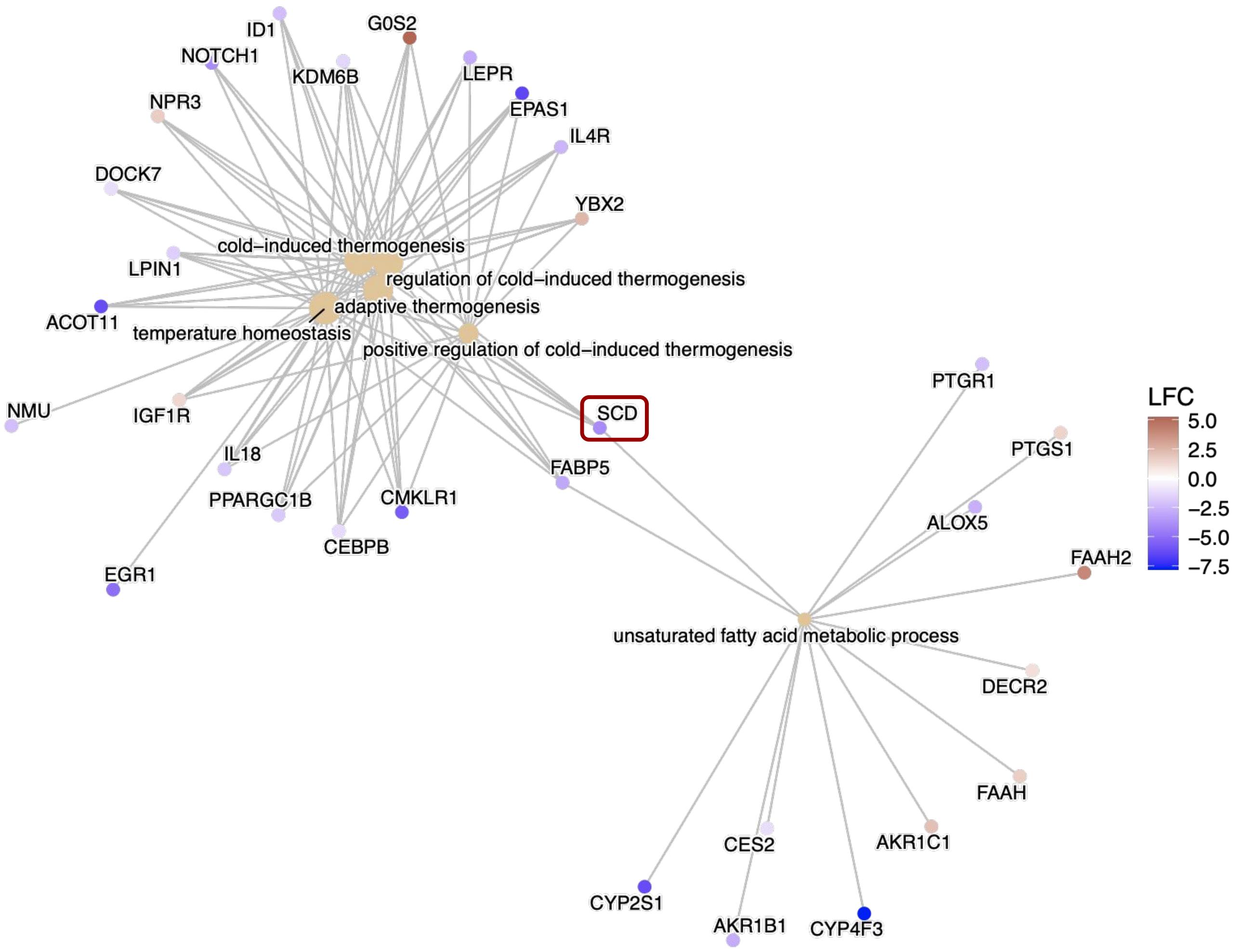
ID1
G0S2
NOTCH1
KDM6B
LEPR
NPR3
EPAS1
IL4R
DOCK7
YBX2
cold−induced thermogenesis
LPIN1
regulation of cold−induced thermogenesis
adaptive thermogenesis
ACOT11
temperature homeostasis
positive regulation of cold−induced thermogenesis
PTGR1
NMU
IGF1R
SCD
LFC
5.0
2.5
0.0
−2.5
−5.0
−7.5
PTGS1
IL18
FABP5
PPARGC1B
CMKLR1
ALOX5
CEBPB
FAAH2
EGR1
unsaturated fatty acid metabolic process
DECR2
FAAH
CES2
AKR1C1
CYP2S1
AKR1B1
CYP4F3

### Figure S7

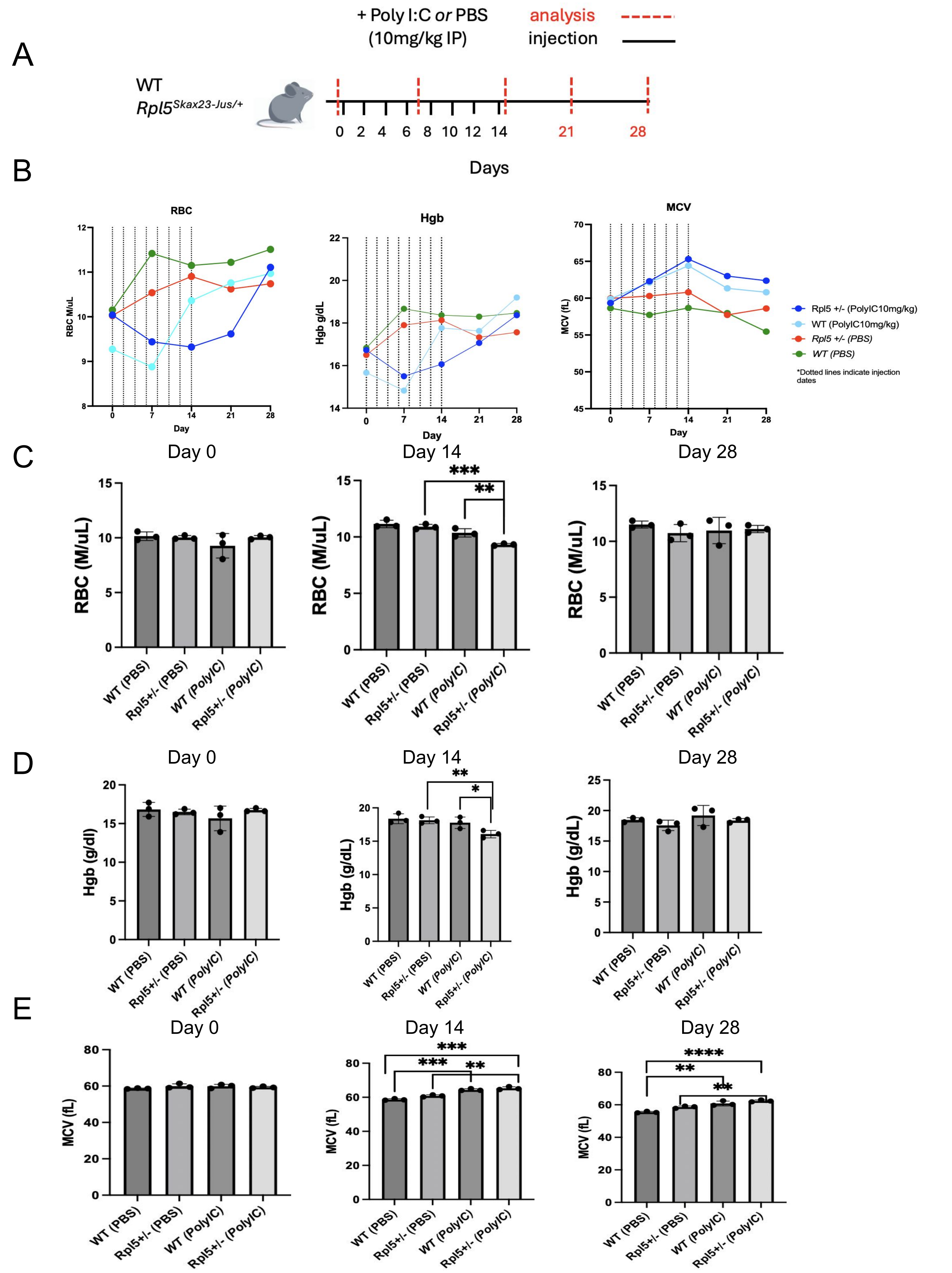
A
+ Poly I:C or PBS
(10mg/kg IP)
analysis
injection
WT
Rpl5Skax23-Jus/+
0 2 4 6 8 10 12 14 21 28
Days
B
RBC
RBC M/uL
Day
Hgb
Hgb g/dL
MCV
MCV (fL)
Rpl5 +/- (PolyIC10mg/kg)
WT (PolyIC10mg/kg)
Rpl5 +/- (PBS)
WT (PBS)
*Dotted lines indicate injection dates
C
Day 0
Day 14
Day 28
RBC (M/uL)
WT (PBS)
Rpl5+/- (PBS)
WT (PolyIC)
Rpl5+/- (PolyIC)
D
Hgb (g/dl)
Hgb (g/dL)
E
MCV (fL)

#### *Figure S8*

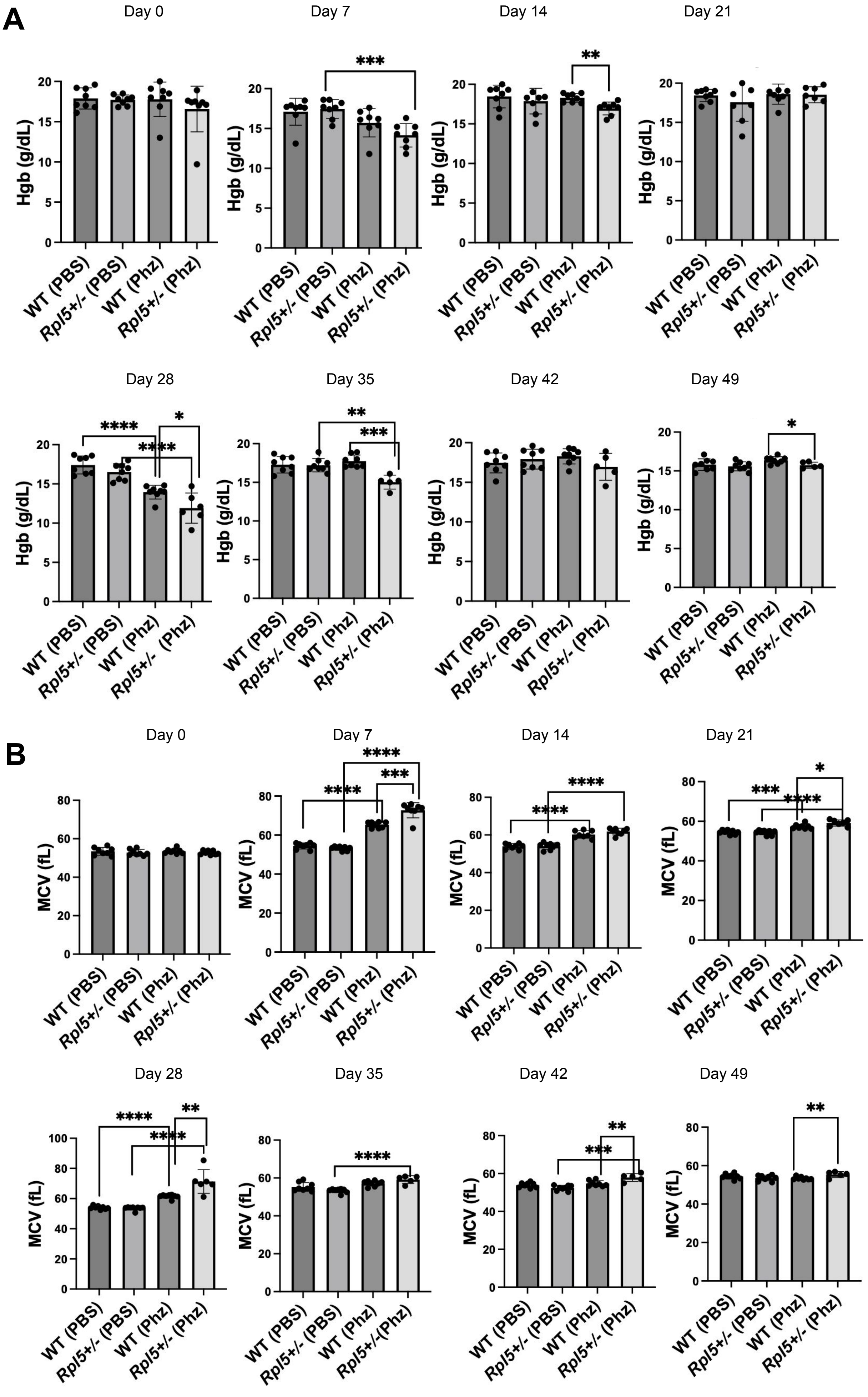
A
Day 0
Day 7
Day 14
Day 21
Day 28
Day 35
Day 42
Day 49
Hgb (g/dL)
WT (PBS)
Rpl5+/- (PBS)
WT (Phz)
Rpl5+/- (Phz)
B
Day 0
Day 7
Day 14
Day 21
Day 28
Day 35
Day 42
Day 49
MCV (fL)
WT (PBS)
Rpl5+/- (PBS)
WT (Phz)
Rpl5+/- (Phz)

*Figure S9*

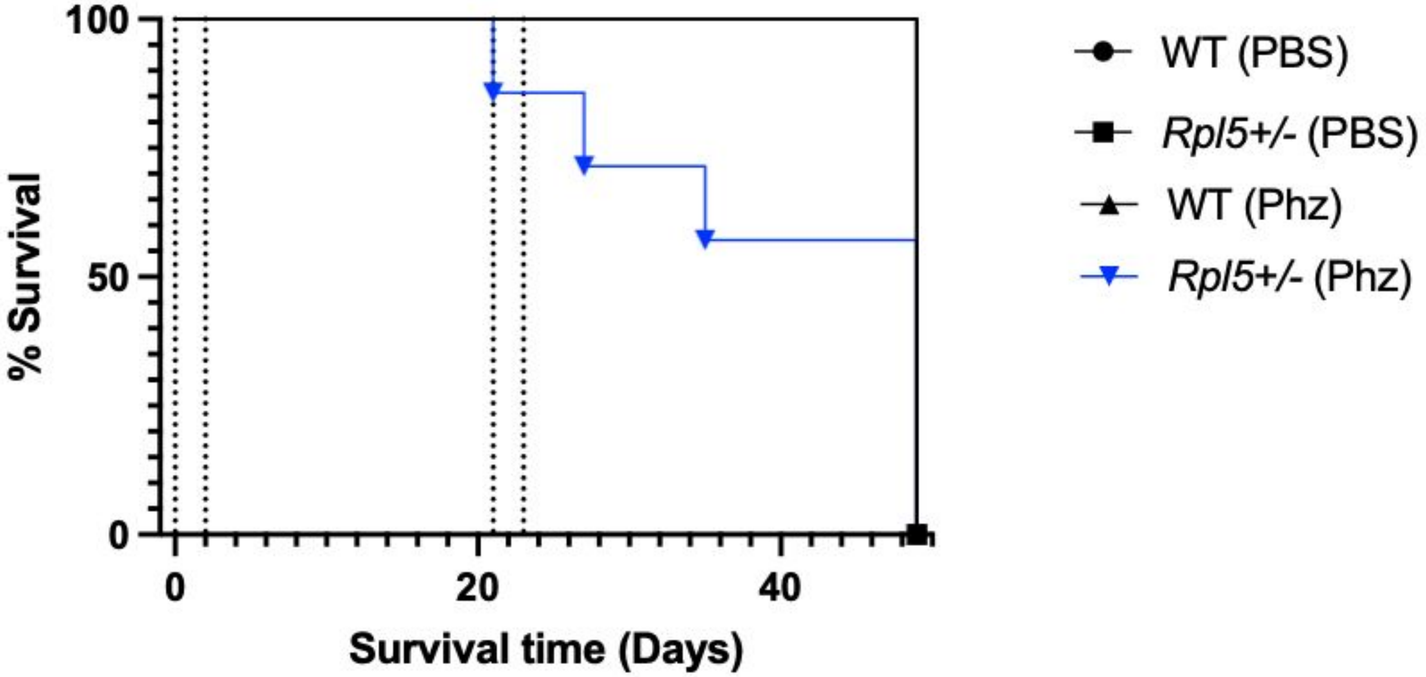

100
50
0
% Survival
0
20
40
Survival time (Days)
WT (PBS)
Rpl5+/- (PBS)
WT (Phz)
Rpl5+/- (Phz)

*Figure S10*

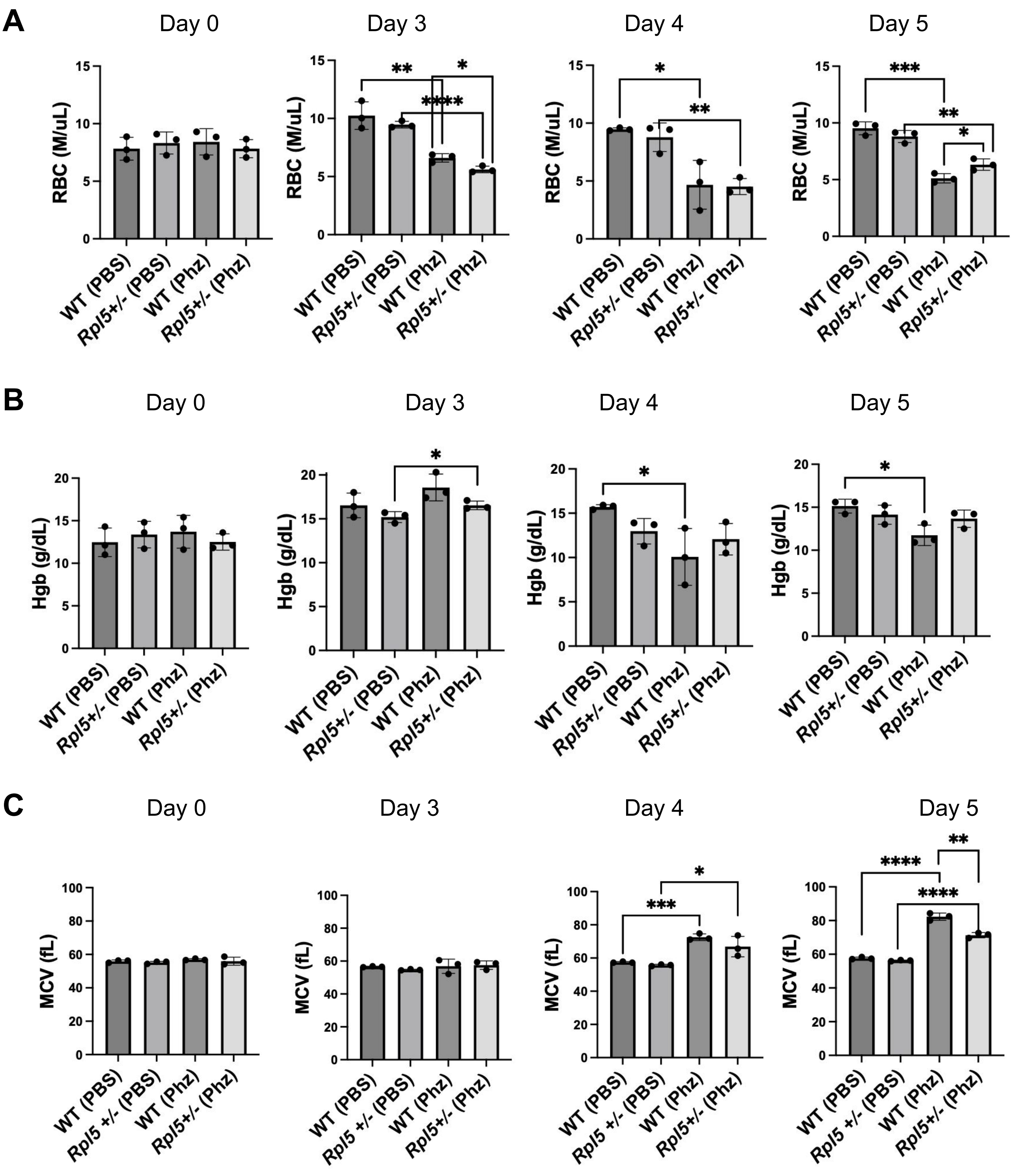
A
Day 0
Day 3
Day 4
Day 5
RBC (M/uL)
WT (PBS)
Rpl5+/- (PBS)
WT (Phz)
Rpl5+/- (Phz)
B
Day 0
Day 3
Day 4
Day 5
Hgb (g/dL)
C
Day 0
Day 3
Day 4
Day 5
MCV (fL)
Rpl5 +/- (PBS)

#### Figure S11

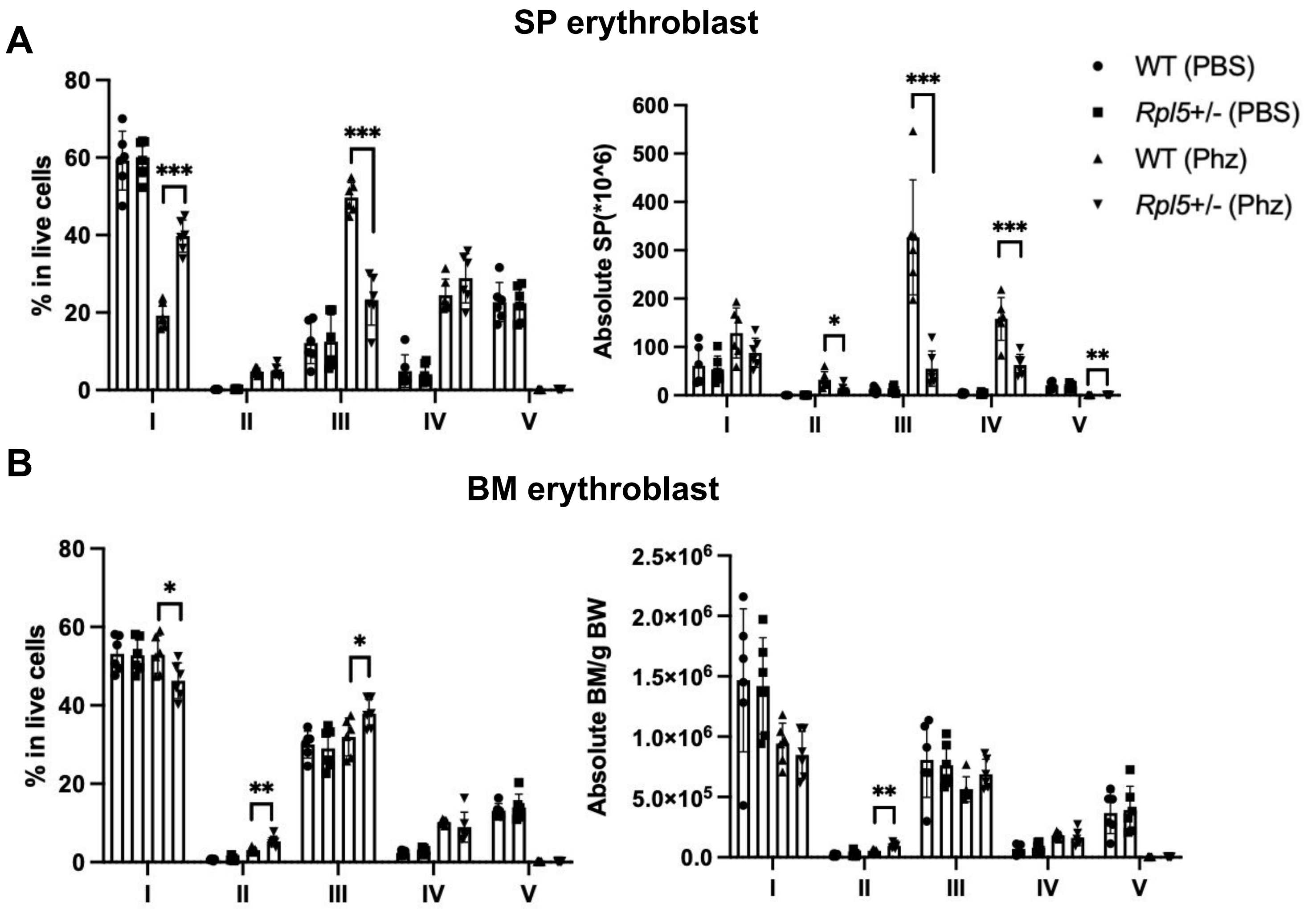
A
SP erythroblast
% in live cells
***
***
I
II
III
IV
V
Absolute SP(*10^6)
***
*
***
**
WT (PBS)
Rpl5+/- (PBS)
WT (Phz)
Rpl5+/- (Phz)
B
BM erythroblast
% in live cells
*
*
**
Absolute BM/g BW
**

### Figure S12

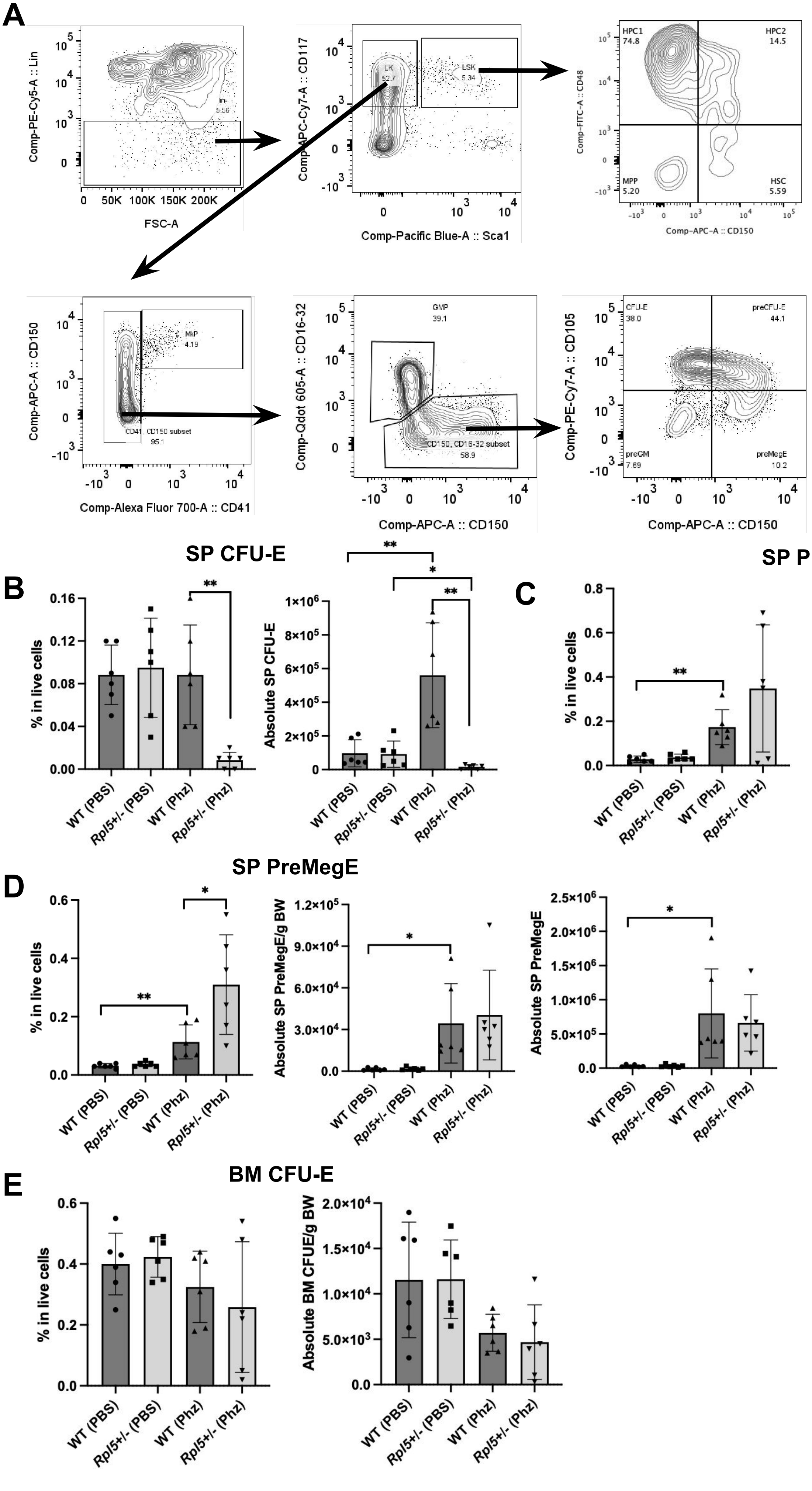

A
Comp-PE-Cy5-A :: Lin
FSC-A
lin-
5.56
Comp-APC-Cy7-A :: CD117
Comp-Pacific Blue-A :: Sca1
LK
52.7
LSK
5.34
Comp-FITC-A :: CD48
Comp-APC-A :: CD150
HPC1
74.8
HPC2
14.5
MPP
5.20
HSC
5.59
Comp-APC-A :: CD150
Comp-Alexa Fluor 700-A :: CD41
MkP
4.19
CD41, CD150 subset
95.1
Comp-Qdot 605-A :: CD16-32
GMP
39.1
CD150, CD16-32 subset
58.9
Comp-PE-Cy7-A :: CD105
CFU-E
38.0
preCFU-E
44.1
preGM
7.69
preMegE
10.2
SP CFU-E
B
% in live cells
Absolute SP CFU-E
SP PreCFU-E
C
% in live cells
Absolute SP PreCFU-E
SP PreMegE
D
% in live cells
Absolute SP PreMegE/g BW
Absolute SP PreMegE
BM CFU-E
E
% in live cells
Absolute BM CFUE/g BW
WT (PBS)
Rpl5+/- (PBS)
WT (Phz)
Rpl5+/- (Phz)

#### *Figure S13*

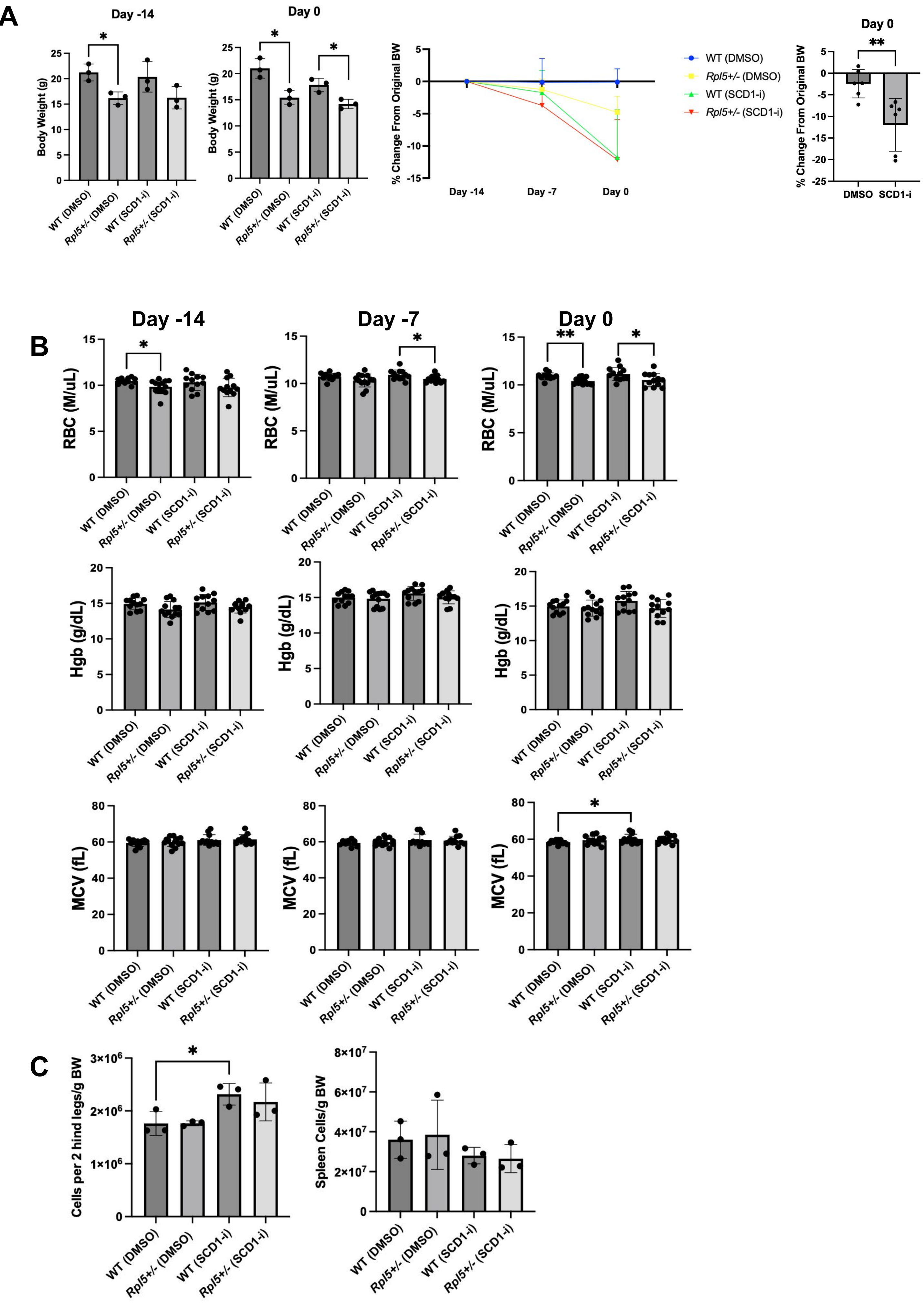

A
Day -14
Day 0
Body Weight (g)
% Change From Original BW
Day -14
Day -7
Day 0
WT (DMSO)
Rpl5+/- (DMSO)
WT (SCD1-i)
Rpl5+/- (SCD1-i)
DMSO
SCD1-i
B
Day -14
Day -7
Day 0
RBC (M/uL)
Hgb (g/dL)
MCV (fL)
C
Cells per 2 hind legs/g BW
Spleen Cells/g BW
