## Supplemental Material for "A lipid metabolism defect is an underlying contributor to Diamond Blackfan anemia syndrome"

### Supplemental Figure Legends

**Table S1: Antibodies used in flow cytometry studies**

**Table S2: Primers used in RT-qPCR studies**

**Table S3: VSD and cleft palate analysis at E15.5** (A) Analysis of E15.5 mice (n=24) demonstrated the presence of VSD in 3/13 *Rpl5*$^{+/-}$ embryos (#3-7, 6-6, 10-2) and cleft palate in 4/13 mutant embryos (#6-1, 6-6, 10-2 and 32-3).

**Figure S1: Summary of the top GO results in different sets of DEGs from the data in Figure 3.** The top 20 significant gene ontology (GO) terms (sorted by the numbers of involved genes) are called from the first experiment's differential expression genes (DEGs). The three panels of plots represent the results from all DEGs, the down-regulated DEGs, and the up-regulated DEGs, respectively.

**Figure S2: The GO results from previously published human data summarizing erythroid and lipid-related terms.** The analyses were performed on the RNA-seq data of a previously published study[39]. Differentially expressed genes (DEGs) were identified by comparing the three control samples and five DBAS samples with an adjusted p-value threshold of 0.05. After that, these DEGs were used for the GO enrichment analysis. (A) shows the dot plots summarizing the significant erythrocyte- and lipid-related terms using all DEGs and down-regulated DEGs, and (B) visualizes the genes involved in these terms.

**Figure S3: The GO results from previously published human data summarizing erythroid and oxidative stress-related terms.** The analyses were performed on the same RNA-seq data of a previously published study[39] as in Figure S2. Differentially expressed genes (DEGs) were identified by comparing the three control and three transfusion-dependent DBAS samples with an adjusted p-value threshold of 0.05. (A) the dot plots summarize the significant erythrocyte- and oxidative stress-related terms using the down-regulated DEGs, and (B) visualizes the genes involved in these terms.

**Figure S4: Summary of the top GO results in different sets of DEGs from the data in Figure 4.** The top 20 significant GO terms (sorted by the numbers of involved genes) from differential expression genes (DEGs) of all three mutant-wild-type comparisons (M-high_WT, M-low_WT and M-all_WT). The three panels of plots represent the results from all DEGs, the down-regulated DEGs, and the up-regulated DEGs, respectively, while each panel compares the results of different mutant-wild-type comparisons labeled on the x-axis.

**Figure S5: The GO term results for data in Figure 4 using the nine specific terms of interest.** (A) Dot plot indicating the terms' significant levels and the numbers of genes for the three mutant-wild-type comparisons when considering all significant genes and down-regulated genes only. Dots with FDR larger than 0.25 are not plotted. Triangular dots mean not significant (FDR ≥ 0.05). (B) The networks related to erythrocyte development, lipid metabolism, and oxidative stress, where the differentially expressed genes (DEGs) are significantly enriched. The points of genes are colored by their log2-fold-change values.

**Figure S6: The GO results from previously published human data summarizing the SCD-involved terms.** The analyses were performed on the same RNA-seq data of a previously published study[39] as Figures S2 and S3. Differentially expressed genes (DEGs) were identified by comparing the three control samples with (A) the five DBAS samples and (B) three transfusion-dependent DBAS samples, respectively. The thresholds of adjusted p-values and the log-2 fold changes were set the same as the numbers in the mice data analyses. (A) and (B) are the networks visualizing the SCD-involved terms, as well as the other DEGs related to these terms. The nodes of SCD were highlighted.

**Figure S7: *Rpl5*$^{+/-}$ mice show delayed erythroid recovery after Poly(I:C) treatment.** (A) Adult female mice were injected with Poly(I:C) at 10mg/kg or PBS by IP. The injection was given every other day starting from Day 0 for two weeks. Peripheral blood was analyzed weekly from Days 0-28. (B) Summary of the weekly plots of RBC, Hgb and MCV. *Rpl5*$^{+/-}$ mice showed significantly lower RBC counts (C) and Hgb levels (D) at Day 14 after Poly(I:C) injection compared to WT mice, with a delay in recovery to normal. (E) Mice injected with Poly(I:C) showed significantly increased MCV level at Days 14 and 28.

**Figure S8: *Rpl5*$^{+/-}$ Phz-treated mice had more significant macrocytic anemia** with (A) significantly lower Hgb and (B) elevated MCV after both Phz treatments. WT and *Rpl5*$^{+/-}$ both showed recovery by days 21 and 49. However, *Rpl5*$^{+/-}$ mice experienced slower recovery of HB and MCV after the second Phz treatment.

**Figure S9: Survival after phenylhydrazine.** Adult mice (n=8 for each group:, 50/50 M:F ratio) were injected with phenylhydrazine (Phz) (60μg/g IP) or PBS on days 0, 2, 21 and 23 (indicated by dotted lines). Death occurred in 3 of 4 *Rpl5*$^{+/-}$ male mice.

**Figure S10**: **Phenylhydrazine time course.** Adult (2 month old) mice were injected with phenylhydrazine (Phz) (60μg/g IP) or PBS on days 0 and 2, and blood counts were obtained on days 0,3,4,5, (n=3 for each group, females). The nadir of the RBC counts was found to be on day 4.

**Figure S11: *Rpl5*$^{+/-}$ mice show altered erythropoiesis after phenylhydrazine treatment**. (A) Quantification of erythroid cells at various stages of differentiation (stages I-V) in spleens and (B) bone marrows. Gating strategy is shown in Figure 5D.

**Figure S12: *Rpl5*$^{+/-}$ mice show reduced spleen CFU-E after phenylhydrazine treatment.** (A) Representative flow cytometry plots showing gating strategies of hematopoietic stem and progenitor populations. Quantification of spleen CFU-E (B) PreCFU-E (C) and PreMegE (D) from WT and *Rpl5*$^{+/-}$ mice with or without Phz treatment. (E) Quantification of bone marrow CFU-E.

**Figure S13: Treatment with SCD1 inhibitor.** Adult mice were pretreated with SCD1 inhibitor (5mg/kg) for 2 weeks and then given Phenylhydrazine on days 0 and day 2. (A) Weights and (B) blood counts were measured at baseline (day -14) and then weekly. There was some weight loss in the SCD1-i treated mice but no blood count change during the pretreatment phase. (C) Quantification of bone marrow and spleen cellularity at day 4.

### Supplemental Methods

#### *Additional mouse procedures*

Peripheral blood was drawn from the lateral saphenous vein into EDTA tubes and blood counts measured with a Drew Scientific Hemavet 950FS. SCD1 inhibitor (CAY10566) stock was dissolved in DMSO (10μg/μL) and diluted (1:10) in PBS 30%

2-Hydroxypropyl-β-cyclodextrin (Sigma Aldrich H107-5G). SCD1 inhibitor treatment was administered daily via oral gavage in 1.25-5 mg/kg doses. Mice treated with SCD1 inhibitor developed blepharoconjunctivitis and were treated with a topical analgesic and an antibiotic. Phenylhydrazine (Sigma Aldrich 114715-5g) was dissolved in PBS (10 mg/mL solution) and administered in 60mg/kg doses. Poly(I:C) HMW VacciGrade (InvivoGen vac-pic) was dissolved in endotoxin-free physiological water and administered in 10 mg/kg doses. Both phenylhydrazine and Poly(I:C) were administered via intraperitoneal injection.

*RT-qPCR*

Total RNA was extracted using RNeasy® Micro Kit (Qiagen, Cat 74004). cDNA synthesis was done by using an iScript cDNA synthesis kit (1708890, Bio-Rad). Quantitative polymerase chain reaction (qRT-PCR) was performed using Power SYBR Green MasterMix (ABI4367659) on an ABI QuantStudio 3 (Thermo Fisher Scientific). Sequences of all primers used in this study are listed in Supplemental 2.

*Flow cytometry*

Mouse bone marrow or spleen cells were harvested and washed with PBS supplemented with 3% FBS. For assessment of erythroid cell differentiation, cells were stained with anti-mouse CD71, Ter119, and CD44 antibodies at 4°C for 30 min in the dark. For HSPC detection, 20 million cells were stained with ZombieAqua, followed by lineage cocktail staining and a combination of antibodies as previously described[1–3] (Supplemental Table 1). Cells were filtered with a 40 µm cell strainer to remove clumps before analysis. Flow analysis was performed using a Fortessa instrument (BD Biosciences) and analyzed on Flowjo (v10.10.0.exe).

*Minigene assay and splice disruption calling*

The 5'UTR (142bp), the first exon (3bp), and the downstream introns (42bp) of *Rpl5* were amplified from the genome of WT and *Rpl5*$^{+/-}$ (c.3+6T>C) mice with overhang primer set:

Forward: 5'-CTTTTGCAAAAAGCTTCAGCCACTCTTTCTCACGTCGCTTG-3'; Reverse: 5'-CATATGCTTTAGCATCTCCGAGCGGGCATCCGCGAGAG-3'.

A *Rpl5* minigene construct was then prepared by cloning these two PCR products, individually, into the pSPL3 vector (Invitrogen, Carlsbad, CA) near HindIII and EcoRV restriction enzyme cutting sites by HiFi assembly (New England Biolabs, Ipswich, MA). The resulting vector was designed to replace the synthetic first exon and preserve the synthetic last exon in the construct. The two minigene constructs were then transiently transfected into 293T cells using FuGENE HD Transfection Reagent (Promega), respectively. Total RNA was harvested at 36 hours post-transfection using RNeasy® Micro Kit, and 5 µg of total RNA was reverse transcribed using SuperScript III First-Strand Synthesis kit (Invitrogen) with oligo(dT) 20 primer following the manufacturer's protocol. Afterwards, as previously described[4,5], the spliced transcript was amplified via semi-nested PCR using outer primer pairs: *Rpl5* Forward 1 (5′-CAGCCACTCTTTCTCACGTC-3′) and JKlab230 reverse (5′-ATCTCAGTGGTATTTGTGAGC-3′), and then inner primer pairs: *Rpl5* Forward 2 (5′-GCCGACTCTGCAGGTCTG-3′) and *Jklab230* reverse. PCR products were purified by SPRIselect Bead-Based Reagent (Beckman Coulter, Brea, CA). Indexed Illumina sequencing adaptors were added by PCR, and the resulting RNA-seq libraries were submitted for paired-end 150-bp sequencing on Illumina HiSeq or NovaSeq instruments. The paired, splice-informative reads were aligned to the reference minigene sequence with the splice-aware read aligner STAR[6]. Reads mapped to the forward strand were kept for further analyses and visualizations (samtools view -F 16). A Python tool, trackplot, was used to visualize the splice reads and disruptive patterns[7].

*Histological analysis, immunohistochemistry, and Terminal deoxynucleotidyl transferase dUTP nick end labeling (TUNEL) staining*

Mouse embryos at embryonic day E14.5 and E15.5 were harvested and fixed with 4% paraformaldehyde (PFA) overnight at 4°C. Tissues were transferred to 30% sucrose in PBS until the tissues sank to the plate bottom. After that, the tissues were embedded in an optimal cutting temperature compound (Thermo Fisher Scientific, Cat #23-730-571), and 10-μm cryosections were prepared by cryostat (Leica CM1850). Hematoxylin and eosin (H&E) staining, immunohistochemistry, and TUNEL staining were done as described previously[8–10]. The primary antibody used was rabbit anti-mouse Phospho-Histone H3 (Ser10) (1:200 dilution, Invitrogen, 44-1190G). TUNEL staining was done using the Click-it Plus TUNEL assay kit (ThermoFisherSci, C10617).

*Bioinformatics analysis*

Intron retention events (IR) were detected with iREAD[11] following its recommended pipeline: https://github.com/genemine/iread. The required annotations of independent introns were retrieved from the GTF file of ENSEMBL GRCm38 assembly release 102 using GTFtools. Raw read counts and FPKM normalized counts were collected for downstream analysis and visualization. Differential IR events were detected by DESeq2[12]. *Rpl5* gene structure was visualized by ggbio[13] using the information from mm10 RefSeq annotation (mm10.ncbiRefSeq.gtf.gz), which was retrieved from: https://hgdownload.soe.ucsc.edu/goldenPath/mm10/bigZips/genes/. Differential gene expression analysis was performed using DESeq2 in R on the gene raw counts estimated in the processing steps. Genes with adjusted p-values ≤ 0.005 and absolute log2 fold changes ≥ 1.0 were considered significant. Heatmaps that visualized the gene expression profiles were plotted by the ComplexHeatmap[14] package, using the normalized counts from DESeq2, which were transformed by the variance stabilizing transformation. Gene-ontology (GO) enrichment analyses, including over-representation analysis (ORA) and gene set enrichment analysis (GSEA), were implemented with clusterProfiler[15] in R. For ORA, only the significant genes with valid Entrez IDs were considered. For GSEA, we excluded the genes without valid adjusted p-values and Entrez IDs and used all of the rest as the background. The annotation databases for querying were *org.Mm.eg.db* and the Mouse MSigDB Collection[16].
